# Integration of microbial and chemical synthesis for the efficient production of plitidepsin, a promising anticancer and antiviral agent

**DOI:** 10.1101/2023.04.24.537896

**Authors:** Haili Zhang, Zhen Hui, Mingwei Cai, Shipeng Huang, Wenguang Shi, Mengdi Liang, Yang Lin, Jie Shen, Minghao Sui, Xuyang Li, Qiliang Lai, Jie Dou, Yun Ge, Min Zheng, Zongze Shao, Xiaozhou Luo, Xiaoyu Tang

## Abstract

Plitidepsin, a marine-derived anticancer medicine, is being tested in phase III clinical trials for treating COVID-19. However, the current supply of plitidepsin relies on laborious chemical synthesis processes. Here, we present a new approach that combines microbial and chemical synthesis to produce plitidepsin. We screened a Tistrella strain library to identify a high-yield didemnin B producer, and then introduced a second copy of the didemnin biosynthetic gene cluster into its genome, resulting in the highest yield of didemnin B reported in the literature. Next, we developed two straightforward chemical strategies to convert didemnin B to plitidepsin, one of which involved a one-step synthetic route giving over 90% overall yield. We also synthesized two new didemnin analogues and assessed their anticancer and antiviral activities. Our findings offer a practical and sustainable solution for producing plitidepsin and its derivatives, potentially expediting didemnin drug development.

---

As of the end of March 2023, the severe acute respiratory syndrome coronavirus 2 (SARS-CoV-2) has infected over 750 million people and caused more than 6.8 million deaths worldwide, according to data from the World Health Organization (WHO) website. The magnitude of the coronavirus disease 2019 (COVID-19) has led to the repurposing of old drugs as antiviral agents for the treatment of the disease^[1]^. Plitidepsin (**1**; Figure 1; also known as dehydrodidemnin B), a drug originally used to treat multiple myeloma, has recently gained great attention as an anti-COVID-19 agent in phase III clinical trials^[2]^. Plitidepsin targets the host eukaryotic translation elongation factor 1A (eEF1A)^[3]^, which is a protein subunit of the eukaryotic translation elongation 1 complex (eEF1) with a canonical role in translational elongation by ribosomes^[4]^. Additionally, eEF1A is involved in other cellular processes such as protein degradation, cellular apoptosis, and actin organization^[5]^. A growing body of evidence shows that eEF1A is hijacked by viruses (especially RNA viruses) during replication, making it an intriguing target for antiviral drug development^[6]^. The unique mode of action confers a broad-spectrum antiviral activity on plitidepsin, making it an ideal candidate to address the variability of viruses such as SARS-CoV-2^[7]^. Moreover, an *in silico* study predicted that didemnin B (**2**; Figure 1), a close analogue of plitidepsin, may have the ability to bind the main protease (Mpro) of coronaviruses^[8]^, which is an attractive drug target against SARS-CoV-2^[9]^.

**Figure 1.**
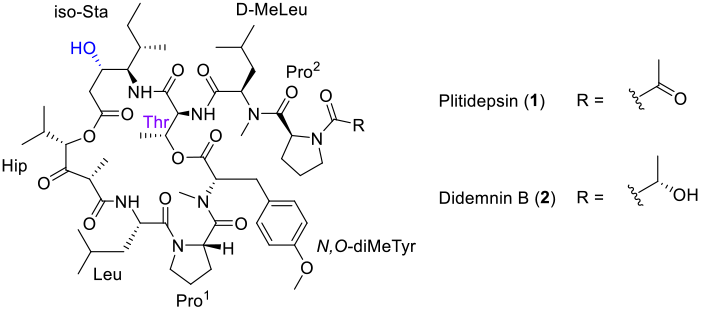
Structures of plitidepsin (**1**) and didemnin B (**2**).

Plitidepsin (**1**, Figure 1) is a member of the didemnin family of natural products, which are cyclic depsipeptides extracted from numerous marine tunicates^[10]^. These compounds have demonstrated excellent antitumor, antiviral, and immunosuppressive properties. Plitidepsin (**1**) was originally isolated from *Aplidium albicans*, an ascidian species obtained in the mediterranean coast of Spain, which was found in small quantities in marine tunicate extracts. The process by which plitidepsin is produced in the natural environment remains unclear, and as a result, the current supply of didemnin-based medicines is dependent on total synthesis, which can involve more than 20 steps and requires many protecting groups and expensive reagents^[11]^. This makes it expensive to use didemnins in research and clinical applications.

Didemnin B (**2**, Figure 1), a member of the didemnin family, was initially discovered in the Caribbean tunicate *Trididemnum solidum* and is structurally similar to plitidepsin^[10a]^. The only difference between the two compounds is a single oxidative state on the lactic side chain (Figure 1). Didemnin B (**2**) was the first marine natural product to be tested in clinical trials as an anticancer agent. Unfortunately, the clinical trials had to be halted due to the cardiac and neuromuscular toxicities associated with its use^[12]^. While the origin of didemnins in tunicates remains unknown, previous studies have identified two marine *α*-proteobacteria, *Tistrella mobilis* and *T. bauzanensis*, that are capable of producing didemnin B (**2**) and a few other minor analogues^[13]^. Based on whole genome sequencing and bioinformatic analysis, Qian, Moore and co-workers proposed that a large nonribosomal peptide synthetase and polyketide synthase(NRPS-PKS) hybrid biosynthetic gene cluster (BGC) on a plasmid of the *T. mobilis* KA081020-065 (isolated from the red sea) is responsible for the biosynthesis of didemnins^[13b]^. This discovery opens the possibility of producing plitidepsin through bioengineering of didemnin biosynthetic pathway or semi-synthesis from didemnins such as didemnin B (**2**). Our initial attempt to pursue this approach, however, was impeded by the very low yield of didemnin B (**2**) in the reported producing strain *T. mobilis* KA081020-065 (∼0.2 mg/L)^[13b]^.

To achieve our goal of obtaining a better didemnin B producer, we collected *T. mobilis* and *T. bauzanensis* strains from our in-house marine culture collections in the Third Institute of Oceanography in China. In total, we obtained 17 *Tistrella* strains, including 14 *T. mobilis* and 3 *T. bauzanensis* strains (Table S1), from the Marine Culture Collection of China (https://mccc.org.cn/). Among them, 15 strains were isolated from the marine sediments or seawater that we collected from Indian Ocean, South China Sea, East China Sea, Yellow Sea, Bering Sea, and Arctic Ocean (Table S1; Figure S1), except for *T. mobilis* MCCC 1A11766 and *T. bauzanensis* MCCC 1A18574 were isolated from wastewater in Thailand and soil in Italy, respectively. We assigned each strain a designated name from L1 to L17 to avoid confusion (Figure 2; Table S1). All strains were submitted to sequence and bioinformatic analysis. AntiSMASH software analysis showed that the proposed didemnin BGC is located on the chromosome for all *Tistrella* isolates (Figure 2a), which is different from the first reported *Tistrella* genome, in which the BGC is located on a plasmid^[13b]^. The didemnin BGC in strain L16, originally isolated from soil, was found to be much shorter than the didemnin BGCs in other strains, with several opening reading frames missing in the middle of the BGC (Figure 2a). Further cultivation using an *Tistrella* fermentation medium (TFM; Supporting Information) revealed that 16 out of 17 *Tistrella* isolates (except for L16) can produce didemnin B (**2**) as the main product (Figure 2b; Supporting Information), indicating that this feature is common across *Tistrella* species, especially those inhabiting marine biospheres. Although the BGCs for synthesizing didemnin B (**2**) are highly similar (87%–100% in identity among *T. mobilis* and 90%–100% in identity among *T. bauzanensis* except for L16) across genomes and metagenome-assembled genomes (MAGs), the didemnin B (**2**) yields differ among the species, with L17 showing the highest yield (22 mg/L) and some species (i.e., L8, L9, L12, and L14) postponing their didemnin B production until the fifth day (Figure 2b). In addition, we recognized that *T. mobilis* L17 has a great adaptation to a wide range of pH and has a much faster growth rate than other used strains in the TFM (Figure S2). Based on these findings, we decided to use *T. mobilis* L17 for further optimization of didemnin B production.

**Figure 2.**
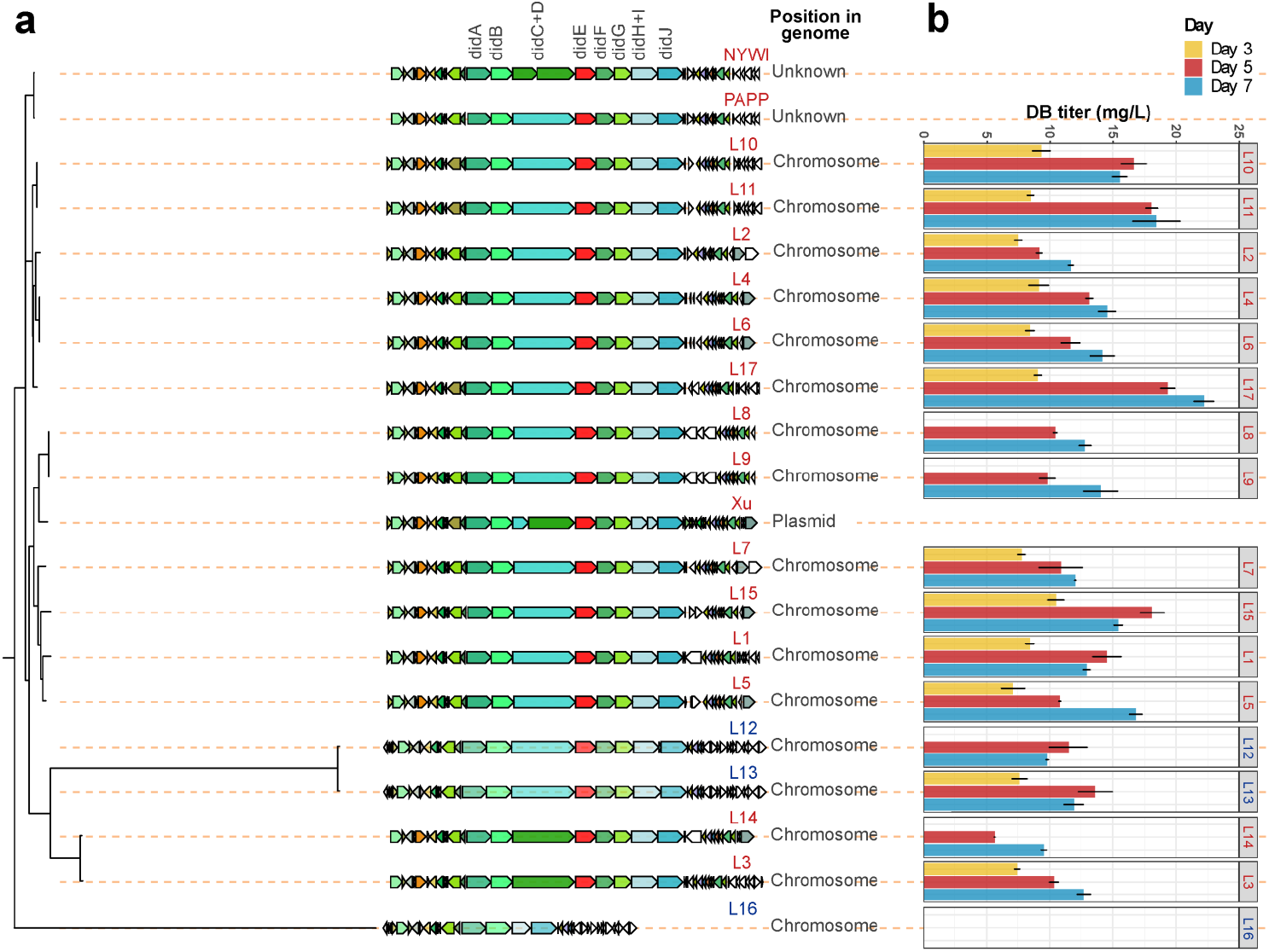
The didemnin BGCs from different *Tistrella* strains (a) and the corresponding didemnin B titers (b) in a fermentation medium. a, CORASON comparison of didemnin BGCs and flanking genes in different *Tistrella* genomes. Didemnin BGCs and flanking genes are predicted by antiSMASH (v6.0.0). Homologous genes are shown using their specific colors. b, Didemnin B titers of *Tistrella* isolates in day 3, 5, and 7. In (a) and (b), *T. mobilis* genomes are marked red, *T. bauzanensis* genomes are highlighted in blue. Error bar means biological triplicate.

Next, we optimized the cultivation conditions by adjusting the fermentation initial pH and seed volume, which resulted in a didemnin B titer of 36 mg/L (Figure S3-S4). To further increase the productivity of the strain, we developed a genetic tool based on sucrose counterselection^[14]^ to manipulate the genome of *T. mobilis* strain (Supporting Information; Table S2-S4; Figure S5). We first tested the method by deleting *didB* (Supporting Information; Table S2-S3; Figure S6), which encodes a NRPS module containing four domains (C-A-KR-T) that is expected to be crucial for didemnin production. The biosynthetic pathway for producing didemnins was only bioinformatically identified in a previous work^[13b]^, so this experiment can provide the direct evidence to support the link between the BGC and didemnin production. As anticipated, the *T. mobilis* L17/*∆ didB(556-826)* mutant strain did not produce any didemnins (Figure S7). This result validates, for the first time, that the proposed BGC is responsible for didemnin production.

The duplication of complete BGCs has been a well-validated approach for improving natural product titers^[15]^. In order to introduce a second copy of the *did* BGC into the genome of the *T. mobilis* L17 strain, a DNA fragment containing the *Φ*C31 phage attachment site *attB* was first inserted into the genome of the *T. mobilis* L17 strain, resulting in the creation of *T. mobilis* L17::*attB* strain (Supporting Information; Table S2-S3; Figure S6). This allows for the introduction of a second copy of the *did* BGC to the genome of the *T. mobilis* L17 strain through *Φ*C31 integrase-mediated integration^[16]^.

To clone the *did* BGC, we conducted PCR screening of a Bacterial Artificial Chromosome (BAC) vector-based DNA library of *T. mobilis* L17. The resulting BAC vector 2-13O, renamed pTZX01, was selected as it is the shortest construct containing the whole *did* BGC (Supporting Information; Table S2-S3; Figure S8). Afterward, we introduced pTZX01 into *T. mobilis* L17::*attB* by triparental intergeneric conjugation with the help of *E. coli* ET12567/pUB307 (Supporting Information). Kanamycin-resistant clones were selected and designated as *T. mobilis* L17::*attB*/pTZX01 (Table S2).

After successfully integrating the second copy of the *did* biosynthetic pathway in *T. mobilis* L17::*attB* strain, we cultured, extracted, and analyzed the supernatant and cellular extracts of the *T. mobilis* L17::*attB*/pTZX01 strain using HPLC and LC-MS. The newly engineered strain was found to be able to produce approximately 60 to 70 mg/L of didemnin B in a 250 mL shake flask with 50 mL liquid medium (Figure S9). To accumulate enough didemnin B, a 10 L culture of the *T. mobilis* L17::*attB*/pTZX01 strain was extracted by ethyl acetate, followed by silica gel column and semi-preparative HPLC separation, resulting in the isolation of approximately 420 mg of didemnin B (**2**) as a white powder (Supporting Information). The structure of the isolated didemnin B (**2**) was confirmed using high-resolution mass spectrometry (HRMS), optical rotation, and extensive NMR spectroscopy (Supporting Information; Table S5-S6; Figure S11-S14; Figure S34) in comparison with previously published data[13b,17].

With enough **2** in hand, we began designing chemical strategies to convert **2** to **1** by oxidizing the lactyl hydroxyl group (Lac-OH) on the side chain. Although **2** bears two secondary hydroxyl groups that are chemically similar, we hypothesized that the hydroxyl group at the *iso*-statine group (*iso*-Sta-OH) is more sterically hindered but electronically more nucleophilic than the Lac-OH. To test this assumption, we attempted to selectively protect each hydroxyl group with different protecting agents. We first used *tert*-butyldimethylsilyl chloride (TBSCl) with imidazole in dimethylformamide (DMF), which predominantly reacted with Lac-OH at 0 ºC after 3 hours, and both hydroxyl groups could be protected after overnight at ambient temperature. This result suggested that the bulky TBS group had better accessibility to the Lac-OH than the *iso*-Sta-OH, as the latter one is adjacent to an *iso*-butyl group. We then used the less bulky protective agent trimethylsilyl group (TMSCl) to react with **2** in DMF at 0 °C for 2 hours, and as expected, TMS preferred to react with the *iso*-Sta-OH, but after prolonging the reaction time, **2** became the predominant product, indicating that TMS protection was labile under the reaction condition. We also observed that *p*-nitrobenzoyl chloride reacted slowly with Lac-OH, while bromomethyl methyl ether did not react with either hydroxyl group. These findings supported our hypothesis that the hydroxyl group at the *iso*-Sta-OH is more sterically hindered but nucleophilic than the Lac-OH. Therefore, we used TMSCl as the protective agent to react with *iso*-Sta-OH to give **3**, which was then oxidized by treatment with Ley-Griffith reagent using *N*-methylmorpholine *N*-oxide (NMO) as a co-oxidant to afford **4** without further purification (Figure 3; Supporting Information). We avoided using Dess–Martin periodinane (DMP) reagent due to its slightly acidic character and potent desilylation. The TMS group was smoothly removed by acetic acid at room temperature to give **1** in 56% overall yield as a white solid (Figure 3; Supporting Information). Because the amide group of the threonine can form a hydrogen bond with either of the carbonyl groups of the pyruvyl unit^[18]^, the two conformers of the compounds **1** and **4** can be readily detected by liquid chromatography–mass spectrometry (LC-MS) with a ratio close to 1:1 (Figure 3). We confirmed the structure of the synthesized compound **1** using high-resolution mass spectrometry (HRMS), optical rotation, and extensive NMR spectroscopy (Supporting Information; Figure S15-S19; Figure S35) in comparison with previously published data^[19]^.

**Figure 3.**
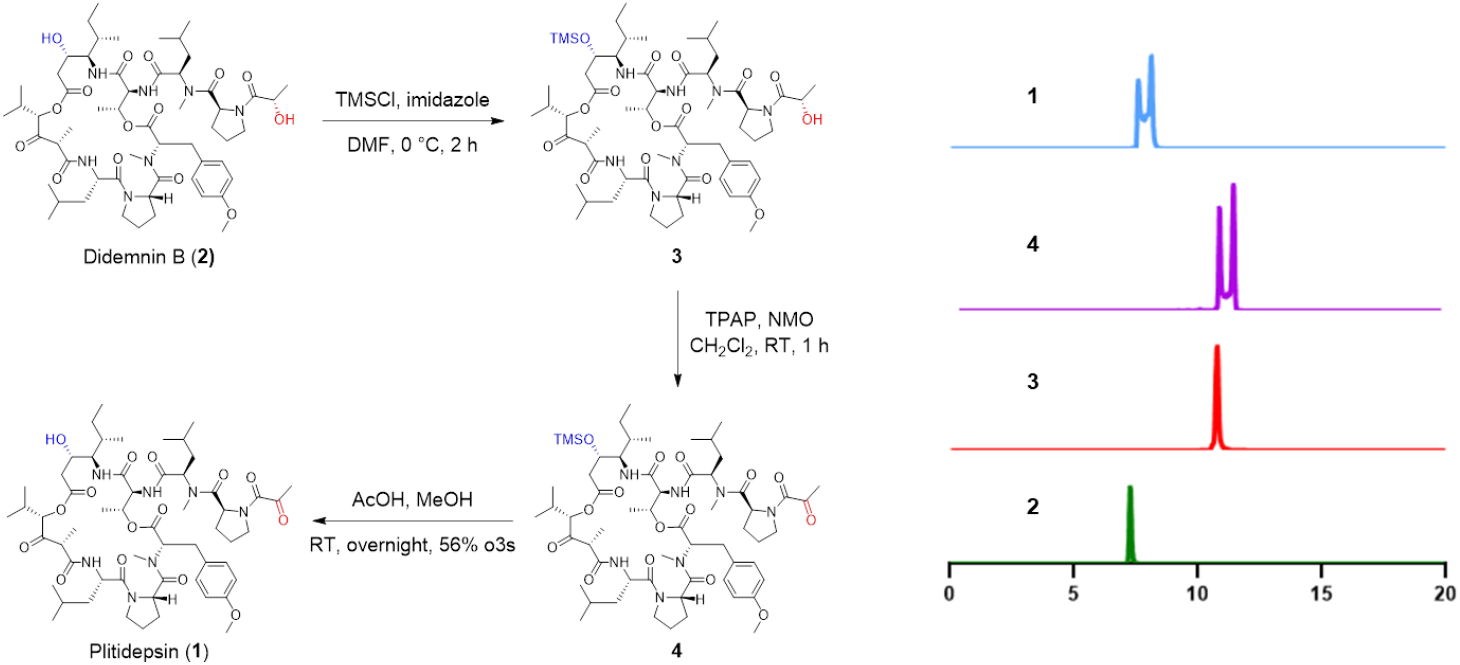
Conversion of didemnin B (**2**) to plitidepsin (**1**) by a TMS protected three-step synthetic route.

Inspired by the successful production of **1** using the protective oxidation strategy, we set out to develop a simpler and more efficient one-step synthetic route. To achieve this, we screened ten different oxidation methods. We showed that the commonly used oxidizing reagent Dess-Martin periodinane (DMP) preferred to oxidize the *iso*-Sta-OH, but lacked the desired selectivity (Figure 4a). We then prepared *tert*-BuDMP to alter the selectivity, but found that its reactivity was significantly reduced, resulting in the detection of only the substrate **2** after 6 hours (Figure 4b). Surprisingly, 2-iodoxybenzoic acid (IBX) proved to be effective at oxidizing the Lac-OH to give **1** in a mixture of DMSO and THF at room temperature. Although a trace amount of the substrate **2** (∼1%) and a small amount of the dual-oxidized byproduct **5** (∼9%) were observed in the reaction mixture, IBX demonstrated selectivity for Lac-OH (Figure 4c). Unfortunately, pyridinium dichromate (PDC) oxidation, modified Pfitzner–Moffatt oxidation, and Ley–Griffith oxidation did not offer selectivity and yielded a mixture of compounds **1, 5**, and **6** under standard conditions (Figure 4d-f). The Parikhing-Doering oxidation displayed relatively low reactivity at room temperature but exhibited good selectivity. This resulted in the major accumulation of compound **1** (77%) with minimal amounts of the byproduct **6** (2%) after overnight incubation (Figure 4g).

**Figure 4.**
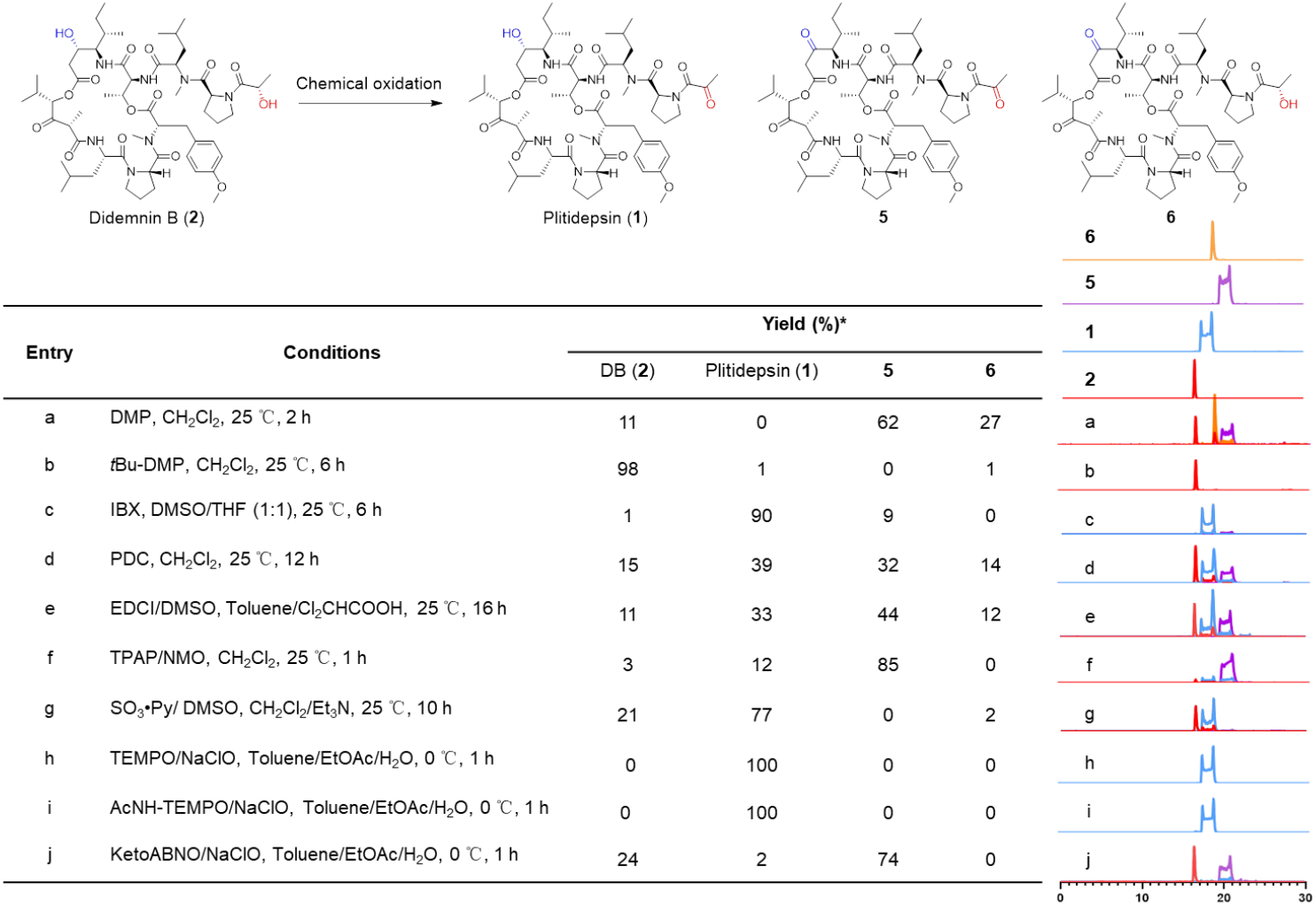
Screening of regioselective oxidants for conversion of didemnin B (**2**) to plitidepsin (**1**). *Yields were calculated by peak area.

Remarkably, the radical catalyst 2,2,6,6-tetramethylpiperidinyloxy (TEMPO), with sodium hypochlorite (NaClO) as the stoichiometric oxidant, yielded only the desired product **1** after one hour at 0 ºC, with no dual-oxidized byproduct detected even after extended reaction times (Figure 4h). Further experimentation with 4-acetamido-TEMPO confirmed the same performance under similar conditions (Figure 4i). There results suggested that the four methyl groups in the TEMPO hinder its ability to attack the *iso*-Sta-OH. In contrast, less sterically hindered 9-azabicyclo[3.3.1]nonan-3-one *N*-oxyl (ketoABNO) was shown to oxidize both hydroxyl groups, forming the dual-oxidized byproduct **5** (Figure 4j). Using TEMPO for oxidation offers several advantages: a) TEMPO oxidation is exclusively reactive for Lac-OH, and no over-oxidation byproduct could be observed even after prolonged incubation time; b) TEMPO is economically cheap and used as a catalyst in the reaction; c) the reaction proceeds very fast, and reaction conditions are mild; d) TEMPO-mediated oxidation is suitable for large-scale oxidation and has been used a variety of industrial applications ^[20]^.

During the screening of the oxidants, the production of the dual-oxidized byproduct **5** and *iso*-Sta-OH oxidized compound **6** was observed. As minor changes in the structure of didemnin B can result in significant differences in bioactivity^[21]^, we attempted to obtain both didemnin analogues to evaluate their biological performances. Compound **5** was efficiently prepared by excess amount of DMP or Ley–Griffith reagent under standard conditions (Supporting Information; Figure S10a; Figure S20-S23; Figure S36). Similar to **1**, compound **5** also exists in a mixture of two conformers in a ratio of 1:1 (Figure S10a). To obtain the *iso*-Sta-OH oxidized product, we first used TBSCl to selectively protect the Lac-OH of **2**, as described above, resulting in **7** (Figure 5b; Supporting Information; Figure S10b; Figure S24-S26; Figure S40). DMP oxidation of **7** led to the formation of **8** in good yield (85%; Supporting Information; Figure S10b; Figure S27-S28; Figure S41). However, attempts to remove the TBS group from **8** in THF were unsuccessful at room temperature using *tetra*-*n*-butylammonium fluoride (TBAF). Additional attempts to deprotect the TBS group were made using both aqueous THF with acetic acid ^[22]^ and aqueous THF with formic acid ^[23]^. The removal of the TBS group from **8** was observed at a very slow rate under the acetic condition. Finally, complete desilylation of **8** was achieved in the formic solution (HCOOH:THF:H2O=3:6:1) to give **6** at room temperature after an overnight incubation in 88% yield (Supporting Information; Figure S10b; Figure S29-S33; Figure S37).

**Figure 5.**
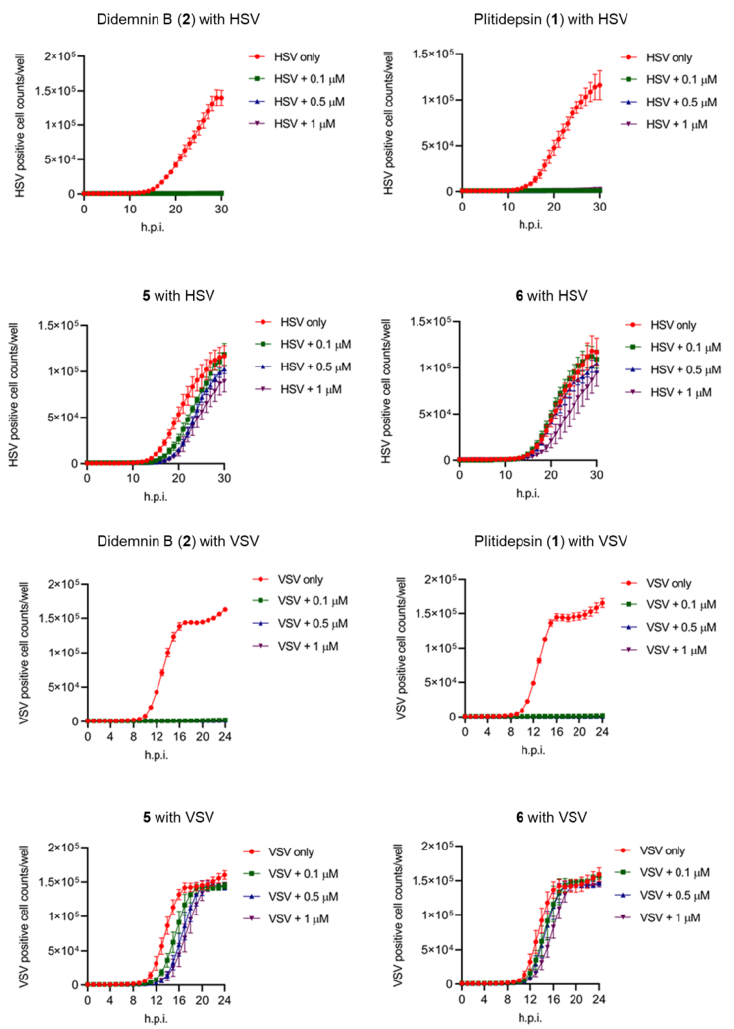
Effects of plitidepsin (**1**) and its derivatives on HSV-1-GFP (0.01 MOI) and VSV-GFP (0.01 MOI) replication in Vero-E6 cells. Real time virus replication in the presence of the indicated doses of didemnin B (**2**), plitidepsin (**1**), **5** or **6** were monitored by IncuCyte S3/SX1. Data are shown as mean ± SEM.

The cytotoxic activities of the purified compounds **1, 2, 5**, and **6** (Figure S42) were tested on the human colon cancer cell line HCT116 and colon carcinoma cell line RKO. As expected, both plitidepsin (**1**) and didemnin B (**2**) showed excellent cytotoxicity against the HCT116 cell line, with an IC50 of around 13 nM and 28 nM (Table S7), respectively. These values were more potent than the control drug docetaxel, whose IC50 was around 69.9 nM. Concerning the RKO cells, all three compounds (**1, 2**, and docetaxel) showed similar potency, with an IC50 of around 20 nM. However, the two new derivatives, **5** and **6**, exhibited strikingly reduced cytotoxicity against both cell lines, displayed IC50 value around 0.5 to 1 µM (Table S7).

Furthermore, the antiviral activities of the four compounds were also tested against both DNA virus and RNA virus. Similarly, both compounds **1** and **2** exhibited excellent antiviral activity against Herpes Simplex Virus (HSV) and Vesicular Stomatitis Virus (VSV) at a concentration of 100 nM (Figure 5; Figure S42-S43). This supports plitidepsin is a broad-spectrum antiviral agent against both DNA and RNA viruses. However, compounds **5** and **6** both showed weak antiviral activities at the concentrations around 1 µM (Figure 5; Figure S43-S44). These observations support the notion that slight modification of didemnin B structure can significantly alter its biological activity. As the *iso*-Sta-OH is crucial to maintain the biological activity of didemnin B, we envision that this position may be used in the future to alter the biological activities of didemnins.

Pervious molecular docking studies suggested that didemnins may be capable of binding to the SARS-COV-2 main protease (3CL protease). Therefore, we conducted further experiments to evaluate the potential of didemnin B (**2**) to inhibit the 3CL protease using a commercially available assay kit. Unfortunately, we did not observe any inhibition (Table S8).

In conclusion, we have developed a new strategy to produce the anticancer and antiviral drug plitidepsin (**1**) and its derivatives, using a combination of microbial synthesis and chemical synthesis to combat the challenges in both aspects. The process utilized an engineered *T. mobilis* strain for sustainable production of didemnin B (**2**), and took advantage of chemical synthesis for efficient conversion of didemnin B (**2**) to plitidepsin (**1**). By optimizing cultural conditions and BGC duplication, we were able to achieve a significant improvement in the titer of didemnin B (**2**), with a yield of 60-70 mg/L, which is the highest titer reported in the literature. Once didemnin B (**2**) was successfully produced, transformation into plitidepsin (**1**) was achieved through two simple synthetic strategies. The first involved selectively protecting the *iso*-Sta-OH and oxidizing the Lac-OH after deprotection to give the final product. The second strategy involved regioselectively oxidizing didemnin B (**2**) to plitidepsin (**1**) in one step with high yield, which was more efficient and cost-effective. The combination of microbial synthesis and chemical synthesis offers a novel, efficient, and economically feasible approach to produce plitidepsin, providing a promising basis for large-scale industrial production.

In addition, we prepared two new didemnin derivatives (**5** and **6**) and evaluated their anticancer and antiviral bioactivities. We observed a significant reduction in both biological activities upon oxidation of *iso*-Sta-OH, indicating the crucial role of *iso*-Sta-OH as a site for further modification. Importantly, our work introduces a genetic tool for manipulating the genome of *T. mobilis*, which will not only facilitate the understanding of the biosynthetic logic of didmnins, but also opens the door for bioengineering of *T. mobilis* to enhance didemnin production and expand its biosynthetic capabilities.

## Acknowledgements

This work was supported by National Natural Science Foundation of China (82173719 to X.T.), Guangdong Province’s Pearl River Recruitment Program of Talents (2021QN02Y855 to X.T.), Guangdong Basic and Applied Basic Research Foundation (2021A1515110334 to M.C. and 2021B1515020049 to X.L.), Shenzhen Bay Laboratory Start-up Funds (21230051 to X.T.), Shenzhen Bay Laboratory Open Program (SZBL2021080601007 to X.T.), and the Shenzhen Bay Scholar Fellowship (to X. L. and X.T.). The authors thank Professor Bradley S. Moore (University of California at San Diego, USA) for providing valuable discussions and the plasmids pCAP01, pJZ001, and pJZ002. We are grateful for the support from mass spectrometry and NMR core facilities in Shenzhen Bay Laboratory.

## Entry for the Table of Contents

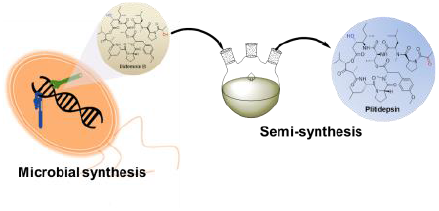

We present a novel approach for the production of plitidepsin, which combines microbial and chemical synthesis. Through genetic engineering of the didemnin B producer *Tistrella mobilis*, we were able to increase the titer of didemnin B and subsequently convert it to plitidepsin via semi-synthesis in a single step. This method provides a sustainable and efficient means of producing plitidepsin.

