## Supporting Information for "Integration of microbial and chemical synthesis for the efficient production of plitidepsin, a promising anticancer and antiviral agent"

Dr. H. Zhang, Z. Hui, Dr. M. Cai, S. Huang, W. Shi, Y. Lin, X. Li, Prof. Dr. Y. Ge, Prof. Dr. X. Tang

Institute of Molecular Chemical Biology, Shenzhen Bay Laboratory, 518132 Shenzhen (China)

S. Huang

Department of Chemistry and Shenzhen Grubbs Institute, Southern University of Science and Technology, 518000 Shenzhen (China)

M. Liang, Prof. Dr. M. Zheng

Institute of Infectious Diseases, Shenzhen Bay Laboratory, 518132 Shenzhen (China)

J. Shen, M. Sui, Prof. Dr. J. Dou

College of Life Science and Technology, China Pharmaceutical University, 211198 Nanjing (China)

Prof. Dr. Q. Lai, Prof. Dr. Z. Shao

Key Laboratory of Marine Genetic Resources, Third Institute of Oceanography, Ministry of Natural Resources, 184 Daxue Road, 361005 Xiamen (China)

Prof. Dr. X. Luo

Center for Synthetic Biochemistry, Shenzhen Institute of Synthetic Biology, Shenzhen Institute of Advanced Technology, Chinese Academy of Sciences,

518055 Shenzhen (China)
[+] These authors contributed equally to this work.

**Table S1** *Tistrella* strain collection from Marine Culture Collection China.

| Designated names | Names | Source | Latitude | Longitude | Location |
|---|---|---|---|---|---|
| L1 | *Tistrella mobilis* MCCC1A01045 | marine sediment | NA | NA | Indian Ocean |
| L2 | *Tistrella mobilis* MCCC1A01199 | marine sediment | 0.13 | -23.62 | Atlantic |
| L3 | *Tistrella mobilis* MCCC1A01369 | marine water | 24.47 | 118.08 | South China Sea |
| L4 | *Tistrella mobilis* MCCC1A01462 | marine water | 6.42 | 95.30 | Indian Ocean |
| L5 | *Tistrella mobilis* MCCC1A02017 | marine water | -31.07 | 58.99 | Indian Ocean |
| L6 | *Tistrella mobilis* MCCC1A02139 | marine water | -26.73 | 66.80 | Indian Ocean |
| L7 | *Tistrella mobilis* MCCC1A02427 | marine water | -7.90 | -16.08 | Atlantic |
| L8 | *Tistrella mobilis* MCCC1A02807 | marine water | 35.18 | 121.67 | Yellow Sea of China |
| L9 | *Tistrella mobilis* MCCC1A02962 | marine water | 36.09 | 120.28 | Yellow Sea of China |
| L10 | *Tistrella mobilis* MCCC1A03759 | marine water | 20.87 | 117.97 | South China Sea |
| L11 | *Tistrella mobilis* MCCC1A03762 | marine water | 20.87 | 117.97 | South China Sea |
| L12 | *Tistrella bauzanensis* MCCC1A07333 | marine water | 53.32 | 169.95 | Arctic Ocean |
| L13 | *Tistrella bauzanensis* MCCC1A07475 | marine water | 67.50 | -169.00 | Arctic Ocean |
| L14 | *Tistrella mobilis* MCCC1A11787 | marine water | NA | NA | South China Sea |
| L15 | *Tistrella mobilis* MCCC1A17629 | marine water | 42.65 | 117.70 | South China Sea |
| L16 | *Tistrella bauzanensis* MCCC1A18574 | soil | 46.52 | 11.35 | Italy |
| L17 | *Tistrella mobilis* MCCC1A11766 | wastewater | 13.74 | 100.52 | Thailand |

**Table S2** Strains and plasmids used in this study.

| Strains | Description | Source |
|---|---|---|
| *E. coli* DH5α | Host for general cloning | Sangon Biotech |
| *E. coli* S17-1 | Donor strain for conjugation | ATCC |
| *E. coli* ET12567/pUB307 | Donor strain for conjugation | ATCC |
| *Tistrella mobilis* L17 | Didemnins producing strain | This study |
| *T. mobilis* L17/Δ*didB556-826* | A *did B* gene mutated strain with the base pairs from 556 to 826 are deleted | This study |
| *T. mobilis* L17::*attB* | A mutant containing the ΦC31 phage attachment site *attB* that is inserted into the gene of *locus_0167* | This study |
| *T. mobilis* L17::*attB*/pTZX01 | A mutant containing a second copy of didemnin biosynthetic gene cluster | This study |
| **Plasmids** | **Description** | **Source** |
| pCAP01 | Plasmid construction, Kanamycin resistance, *oriT*, phiC31, *attP* | [1] |
| pJZ001 | Plasmid construction, Gentamycin resistance, *oriT* | [1] |
| pJZ002 | Plasmid construction, temperature-sensitive *oriC* | [2] |
| pTHZ000 | A plasmid for making gene deletion in *T. mobilis*, Gentamycin resistance, *oriT*, phiC31, *attP*, *sacB* | This study |
| pTHZ001 | A plasmid for making gene deletion in *T. mobilis* with a MCS, Gentamycin resistance, *oriT*, phiC31, *attP*, *sacB* | This study |
| pTHZ001::*didB*(556-826) | pTZ002 derivative for deleting the gene *didB*(556-826) in *T. mobilis* L17 | This study |
| pTHZ001::*0167attB* | pTZ002 derivative for introducing *attB* in *T. mobilis* L17 | This study |
| pTZX001 | pMSBBAC1 derivative containing a 97 kb insert that covers the whole didemnin BGC | This study |

**Table S3** Primers used in this study.

| Name | Sequence |
|---|---|
| JZ002_F | TTTTCTTTTATTGTTTGTTAGTCTTGATGCTTCACTGATAGATACAAGAGCCATAAG |
| JZ002_R | CCCACGACCCCTACGTACAGGCAGGTCTGTTTGCCAGGAACTGCTGAACAGCAAAAAG |
| JZ001_F | ACGTAGGGGTCGTGGGCCACGAAGGCGTGCACGAGTACTCCGCGGCGTTGTGACAAT |
| JZ001_R | AACCTGCCATAGGCCGGCCGAATTGACATAAGCCTGTTCGGTTCG |
| CAP01_F | TTCGGCCGGCCTATGGCAGGTTGGGCGTCGCTTGGTCGGTC |
| CAP01_R | TTCTTCGTCTTGCATGCCTGCAGGTCGACGGATCTTTTCCGCTG |
| hr2800_apali_LF | TGGGCCACGAAGGCGTGCACGATGCGACCGCCTGGGTCGCGGACGACCTG |
| hr2800_hindiii_LR | GGAAGGGCAAAGCTTCAGCCCGTGCAGATCCGGCGCACAGCGGAACAGGC |
| hr2800_hindiii_RF | CGGGCTGAAGCTTTGCCCTTCCGGATCAGCCCCGCGCGGACGGAAGCCCT |
| hr2800_xbai_RR | CTATACTTTCTAGAAGGATGCAGGCCCAGAGTGCCTCGATATGGCGGGAGAGAT |
| hr0167attB_apali_LF | ACGAAGGCGTGCACCAGTGAGGCCCTGAAAGTGGCTGGCATCGTGCGCGATGCCGGTG |
| hr0167attB_hindiii_LR | ATCGAAGCTTTGCGTCTGCTGCGGCGGGCCGGCCGGTGAACGATCGGC |
| hr0167attB_kpni_RF | ATCGGGTACCTCGCCCGGCCCGCCGCCGGCGCAACTTCGCCTGGACGCGGAC |
| hr0167attB_xbai_RR | CTATACTTTCTAGAAATCCGAAAAACAAATTGCATTCATTTATGTCACATCGTC |
| 0167attB_F | AGCTTCGGTGCGGGTGCCAGGGCGTGCCCTTGGGCTCCCCGGGCGCGTACTCCACCGGTAC |
| 0167attB_R | CGGTGGAGTACGCGCCCGGGGAGCCCAAGGGCACGCCCTGGCACCCGCACCGA |
| FDB-F1 | AACACCCCCTCGCAGGTTGC |
| FDB-R2 | AAATCCGGTGGGCCGAAATGCG |
| FDB-F3 | ACCACAGCATGACCAATGC |
| FDB-R4 | ACCGCATGGATATGGTCGCCA |
| FDB-F5 | TGAGCTGGGACGACGACGAAGA |
| FDB-R6 | ATCGGTCTAAAGCCCCGAATG |

**Table S4** The sequence of synthesized *sacB* gene.

| |
|---|
| TGCAGGCATGCAAGACGAAGAATCCATGGGTATGGACAGATCCTCTTTAGgcccgtagtctgcaaatccttttatgattttctatcaaacaaaagaggaaaatagaccagttgcaatccaa<br>acgagagtctaatagaatgaggtcgaaaagtaaatcgcgcgggtttgttactgataaagcaggcaagacctaaaatgtgtaaagggcaaagtgtatactttggcgtcaccccttacatattttaggtctttttttattgtgcgtaact<br>aacttgccatcttcaaacaggagggctggaagaagcagaccgctaacacagtacataaaaaaggagacatgaacgatgaacatcaaaaagtttgcaaaacaagcaacagtattaacctttactaccgcactgctggca<br>ggaggcgcaactcaagcgtttgcgaaagaaacgaaccaaaagccatataaggaaacatacggcatttcccatattacacgccatgatatgctgcaaatccctgaacagcaaaaaaatgaaaaatatcaagttcctgaatt<br>cgattcgtccacaattaaaaatatctcttctgcaaaaggcctggacgtttgggacagctggccattacaaaacgctgacggcactgtcgcaaactatcacggctaccacatcgtctttgcattagccggagatcctaaaaatgc<br>ggatgacacatcgatttacatgttctatcaaaaagtcggcgaaacttctattgacagctggaaaaacgctggccgcgtctttaaagacagcgacaaattcgatgcaaatgattctatcctaaaagaccaaacacaagaatgg<br>tcaggttcagccacatttacatctgacggaaaaatccgtttattctacactgatttctccggtaaacattacggcaaacaaacactgacaactgcacaagttaacgtatcagcatcagacagctctttgaacatcaacggtgtag<br>aggattataaatcaatctttgacggtgacggaaaaacgtatcaaaatgtacagcagttcatcgatgaaggcaactacagctcaggcgacaaccatacgctgagagatcctcactacgtagaagataaaggccacaaata<br>cttagtatttgaagcaaacactggaactgaagatggctaccaaggcgaagaatctttatttaacaaagcatactatggcaaaagcacatcattcttccgtcaagaaagtcaaaaacttctgcaaagcgataaaaaacgcac<br>ggctgagttagcaaacggcgctctcggtatgattgagctaaacgatgattacacactgaaaaaagtgatgaaaccgctgattgcatctaacacagtaacagatgaaattgaacgcgcgaacgtctttaaaatgaacggcaa<br>atggtacctgttcactgactcccgcggatcaaaaatgacgattgacggcattacgtctaacgatatttacatgcttggttatgtttctaattctttaactggcccatacaagccgctgaacaaaactggccttgtgttaaaaatggatc<br>ttgatcctaacgatgtaacctttacttactcacacttcgctgtacctcaagcgaaaggaaacaatgtcgtgattacaagctatatgacaaacagaggattctacgcagacaaacaatcaacgtttgcgccaagcttcctgctgaa<br>catcaaaggcaagaaaacatctgttgtcaaagacagcatccttgaacaaggacaattaacagttaacaaataaaaaCGCAAAAGAAAATGCCGATATCCTATTGGCATTTTCTTTTATTGTT<br>TGTTAGTCTT |

**Table S5** $^{1}H$ NMR (400 MHz, $CDCl_3$) and $^{13}C$ NMR (101 MHz, $CDCl_3$) for didemnin B (**2**).

| | **Position** | **$\delta_C$** | **$\delta_H$ (*J* in Hz)** | | **Position** | **$\delta_C$** | **$\delta_H$ (*J* in Hz)** |
|---|---|---|---|---|---|---|---|
| isoSta | C2Ha | 38.8 | 2.66, m | *N,O*-diMeTyr | CO | 168.5 | |
| | C2Hb | | 3.27, m | | C2H | 66.5 | 3.58, m |
| | C3H | 68.0 | 4.06, m | | C3Ha | 33.9 | 3.18, m |
| | C4H | 55.3 | 4.10, m | | C3Hb | | 3.38, m |
| | C5H | 34.0 | 1.85, m | | C4 | 130.0 | |
| | C5-CH3 | 14.7 | 0.91, m | | C5H | 130.3 | 7.07 (d, 8.1) |
| | C6Ha | 27.1 | 1.43, m | | C6H | 114.1 | 6.85 (d, 8.1) |
| | C6Hb | | 1.20, m | | C7 | 158.6 | |
| | C7H3 | 11.7 | 0.92, m | | OCH3 | 55.3 | 3.80 (s) |
| | NH | | 7.22, d (9.6) | | NCH3 | 38.7 | 2.56 (s) |
| Hip | CO | 169.7 | | Thr | CO | 169.4 | |
| | C2H | 49.5 | 4.25, q (6.7) | | CαH | 57.7 | 4.55, m |
| | C2-CH3 | 15.3 | 1.33, d (6.8) | | CβH | 70.4 | 5.41, m |
| | C3 | 205.0 | | | CγH3 | 16.3 | 1.39, m |
| | C4H | 81.5 | 5.19, d (3.2) | | NH | | 7.66 (d, 5.0) |
| | C5H | 31.3 | 2.36, m | D-MeLeu | CO | 171.7 | |
| | C5-CH3 | 16.9 | 0.88, m | | CαH | 54.9 | 5.38, m |
| | C6H3 | 18.6 | 0.91, m | | CβHa | 36.2 | 1.70, m |
| Leu | CαH | 49.5 | 4.81, m | | CβHb | | 1.84, m |
| | CβHa | 41.3 | 1.22, m | | CγH | 24.9 | 1.43, m |
| | CβHb | | 1.61, m | | CδH3 | 21.4 | 0.87, m |
| | CγH | 24.9 | 1.52, m | | CδH3 | 23.4 | 0.90, m |
| | CδH3 | 20.9 | 0.91, m | | NCH3 | 31.3 | 3.15 (s) |
| | CδH3 | 23.7 | 0.94, m | Pro2 | CO | 172.9 | |
| | NH | | 7.82 (d, 9.3) | | CαH | 56.7 | 4.74, m |
| Pro1 | CO | 170.6 | | | CβHa | 28.4 | 1.98, m |
| | CαH | 57.2 | 4.64, m | | CβHb | | 2.22, m |
| | CβHa | 27.9 | 1.77, m | | CγHa | 26.0 | 1.97, m |
| | CβHb | | 2.14, m | | CγHb | | 2.23, m |
| | CγHa | 25.0 | 2.04, m | | CδHa | 47.0 | 3.57, m |
| | CγHb | | 2.14, m | | CδHb | | 3.68, m |
| | CδHa | 47.0 | 3.61, m | Lac | CO | 173.9 | |
| | CδHb | | 3.70, m | | CαH | 66.0 | 4.39, q (6.6) |
| | | | | | CβH3 | 20.2 | 1.39, m |

**Table S6** Comparison of specific rotation of didemnin B (**2**). Entry i and ii are reported specific rotation from the literatures, and entry iii is the specific rotation for the isolated didemnin B in this study, which was measured on a JASCO P-2000 Polarimeter at 20 °C in $CH_2Cl_2$.

| **Entry** | | **Temperature (°C)** | **Specific Rotation ($CH_2Cl_2$)** |
|---|---|---|---|
| i | didemnin B | 25 | -78 (6.91) [4] |
| ii | didemnin B | 25 | -83 (0.3) [5] |
| iii | didemnin B | 20 | -68 (0.53) |

**Table S7** Cytotoxicity of plitidepsin (1), didemnin B (2), 5 and 6 against cancer cell lines HCT116 and RKO. Docetaxel was used as positive control.

| Cell Line | Name | $IC_{50}$ (nM) | SD |
|---|---|---|---|
| HCT116 | docetaxel | 69.95 | 20.4963 |
| | didemnin B (**2**) | 13.07 | 0.8407 |
| | plitidepsin (**1**) | 28.36 | 3.3509 |
| | **5** | 1027.90 | 126.6749 |
| | **6** | 571.80 | 27.5102 |
| RKO | docetaxel | 16.51 | 11.80 |
| | didemnin B (**2**) | 22.53 | 14.05 |
| | plitidepsin (**1**) | 24.02 | 6.65 |
| | **5** | 1534.37 | 723.90 |
| | **6** | 1113.47 | 322.21 |

**Table S8** The inhibition of didemnin B (2) on the SARS-COV-2 main protease (3CL protease)**.** Ebselen was used as positive control. The Assay Kit was purchased from Beyotime Biotechnology (China) and used followed its instruction.

| Entry | Blank control | 100% enzymatic control | Ebselen (2 μM) | didemnin B (1 μM) | didemnin B (10 μM) | didemnin B (100 μM) |
|---|---|---|---|---|---|---|
| A | 3291 | 50630 | 27122 | 47966 | 50502 | 50809 |
| B | 3167 | 50495 | 26114 | 50038 | 49872 | 49826 |
| C | 2926 | 47244 | 25720 | 49952 | 49952 | 49952 |
| Inhibition ratio | | | | 0.00% | -0.01% | -0.01% |

**Figure S1** Global distribution of the *Tistrella* strains used in this study.

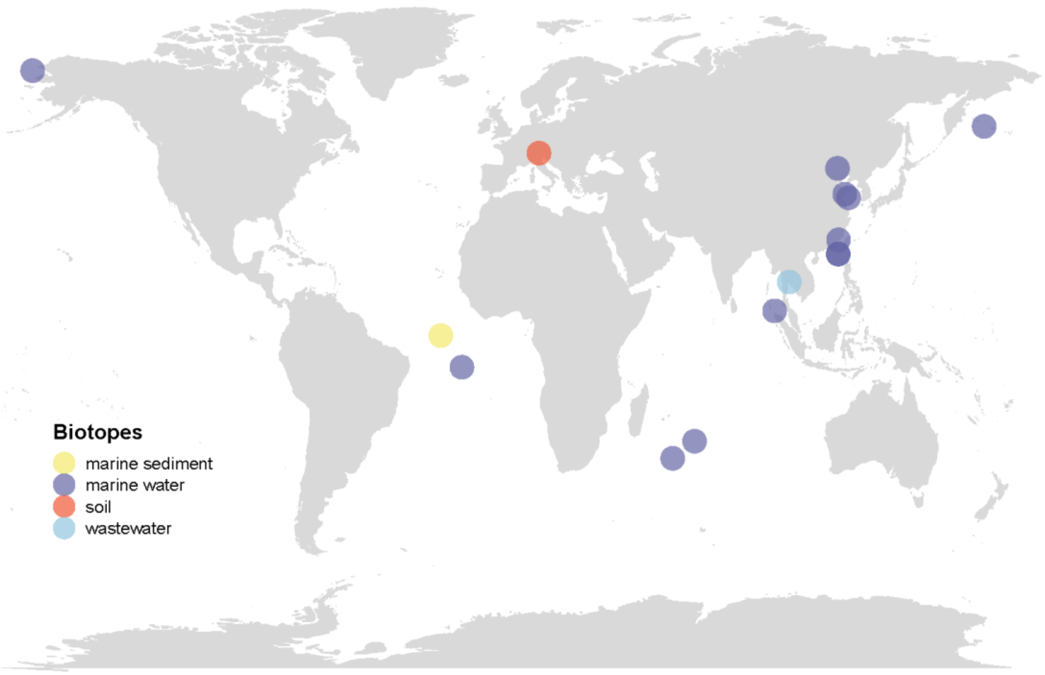

**Figure S2** Growth state of selected *Tistrella* strains in TFM with different initial pH after 24 hours incubation.

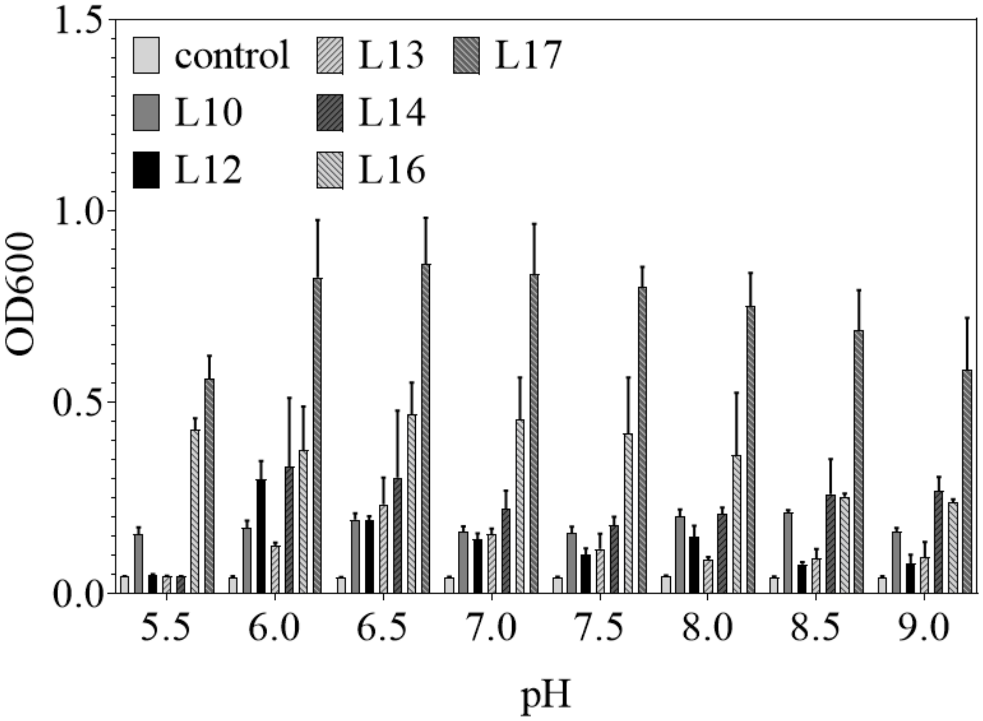

**Figure S3** The titers of didemnin B in the TFM with different initial pH values.

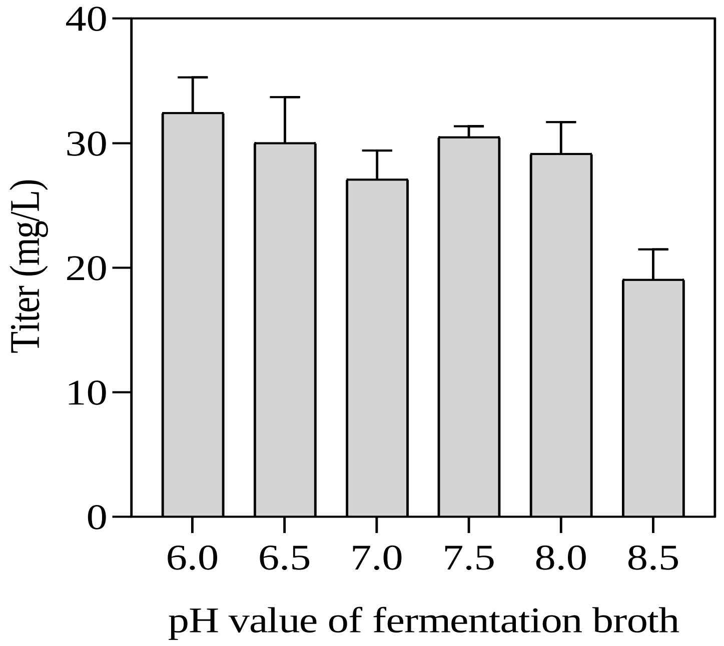

**Figure S4** The titers of didemnin B in the TFM with different seed volumes.

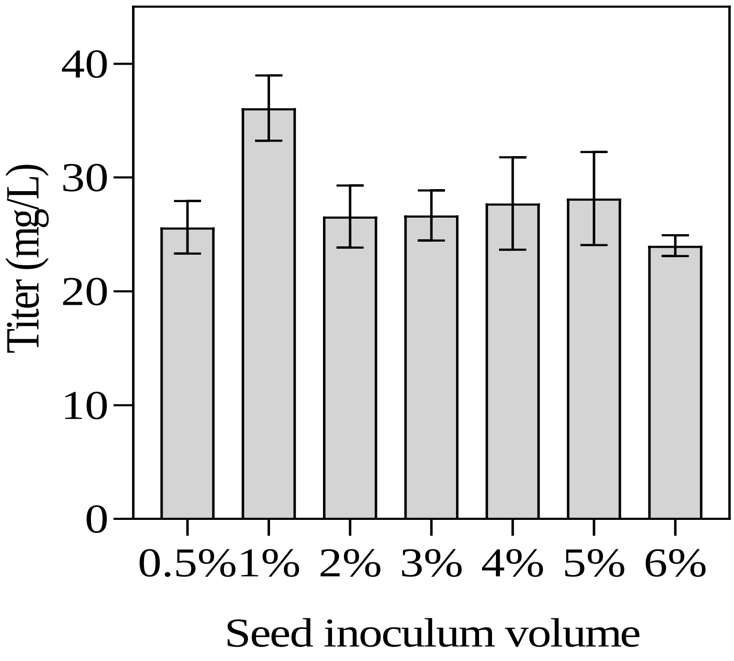

**Figure S5** Plasmid map of pTHZ001. The vector constructed by assembling four different functional elements from pJZ001 (Tan), pJZ002 (Lavender), pCAP01 (Lime), and synthesized sacB component (Red).

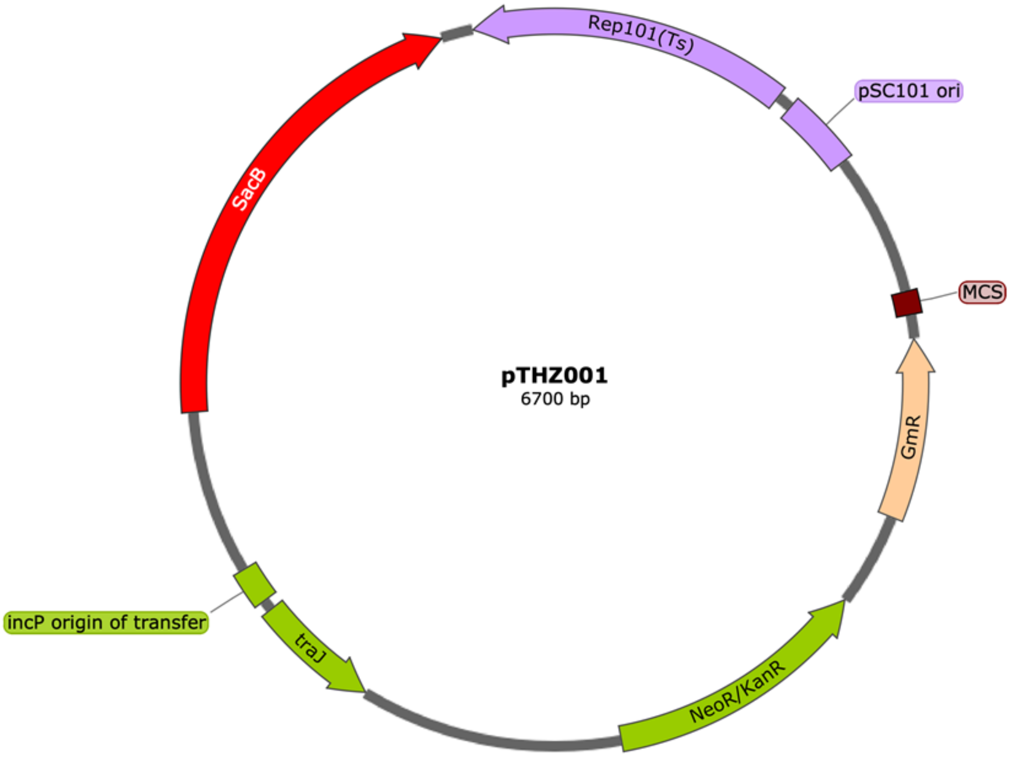

**Figure S6** The Sanger sequencing for verifying *T. mobilis* L17/Δ*didB(556-826)* and *T. mobilis* L17::*attB* mutants. The Upper scheme shows the deletion of *didB* from the base pair number *556 to 826*. The bottom scheme reveals the deletion of locus_*0167* while insertion of *attB* element, which is annotated in green color in the map.

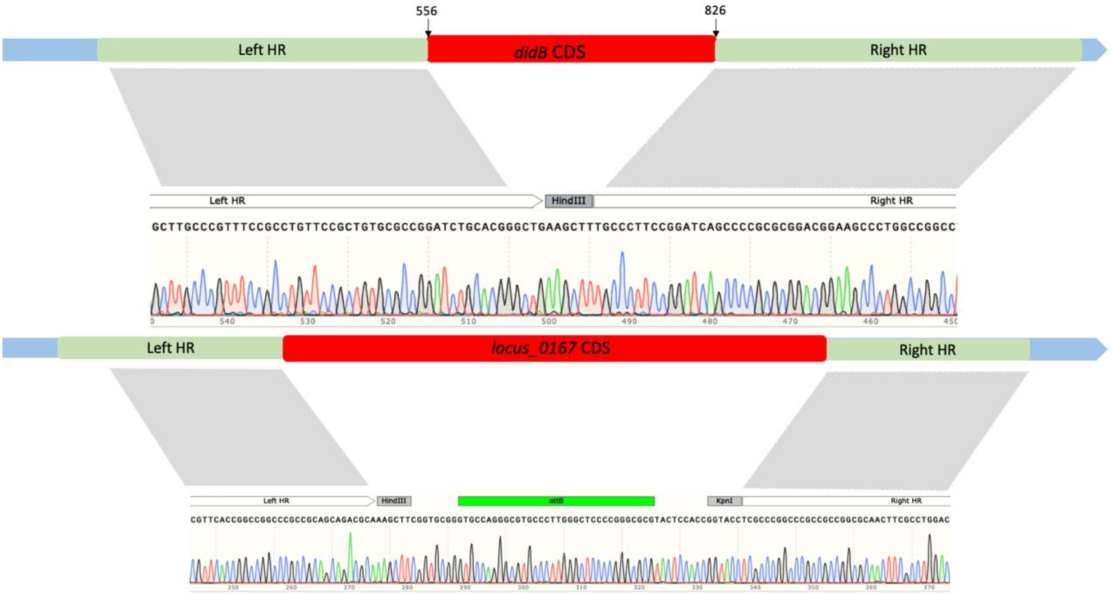

**Figure S7** Comparative LC-HRMS analysis of metabolites from *T. mobilis* L17 and *T. mobilis* L17/Δ*didB*(*556-826*). The abbreviations used are: DB, didemnin B; DX, didemnin X; DY, didemnin Y; NDB, nordidemnin B; NDX, nordidemnin X; NDY, nordidemnin Y; HDB, [Hysp$^2$]didemnin B. 1130 were deduced to be the hydrolytic products of didemnin B dissociated from the two ester bonds.

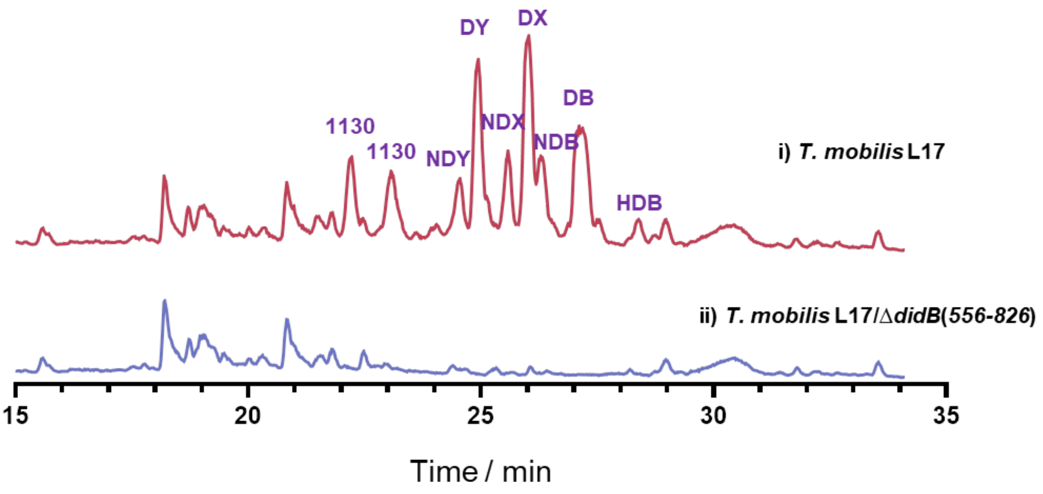

**Figure S8** Screening of a BAC library for vectors carrying didemnin BGC.

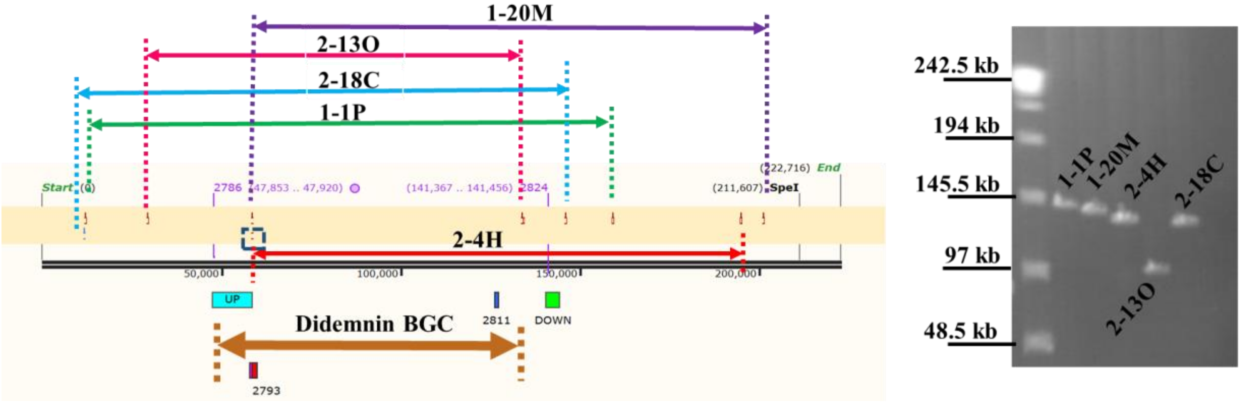

**Figure S9** Growth curve and titers of *T. mobilis* L17::*attB*/pTZX01 in TFM.

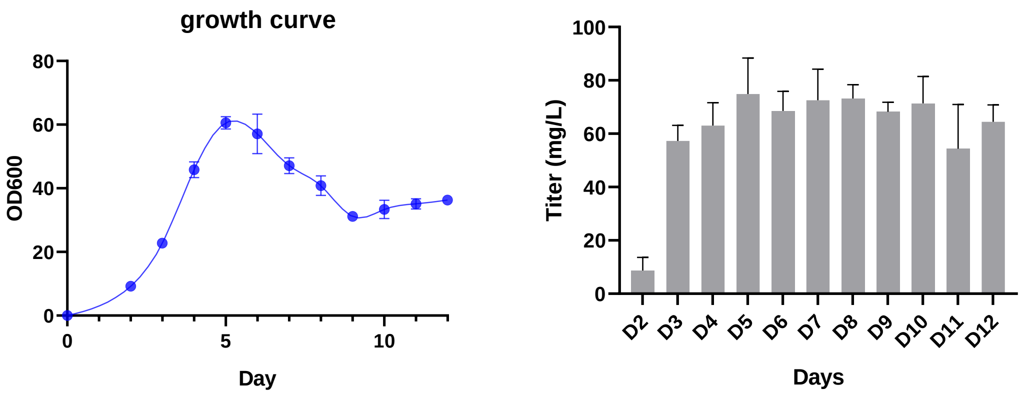

**Figure S10** Semi-synthesis of compounds **5** (a) and **6** (b).

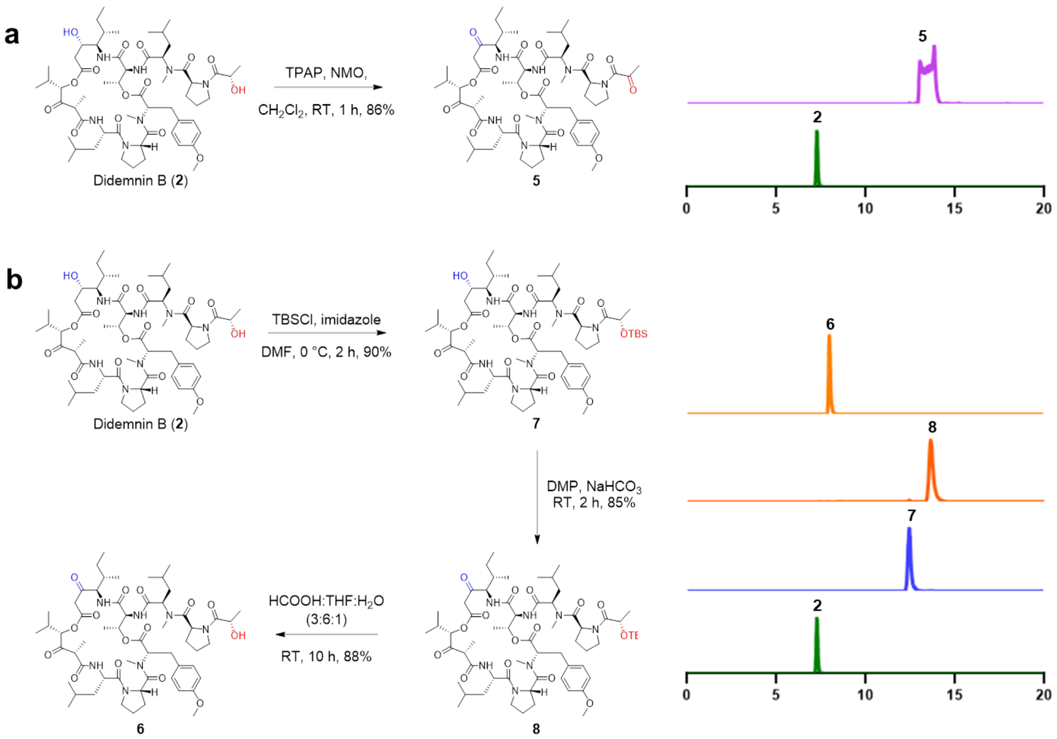

**Figure S11** [1]H NMR spectrum of didemnin B (**2**, 400 MHz). Solvent: $CDCl_3$.

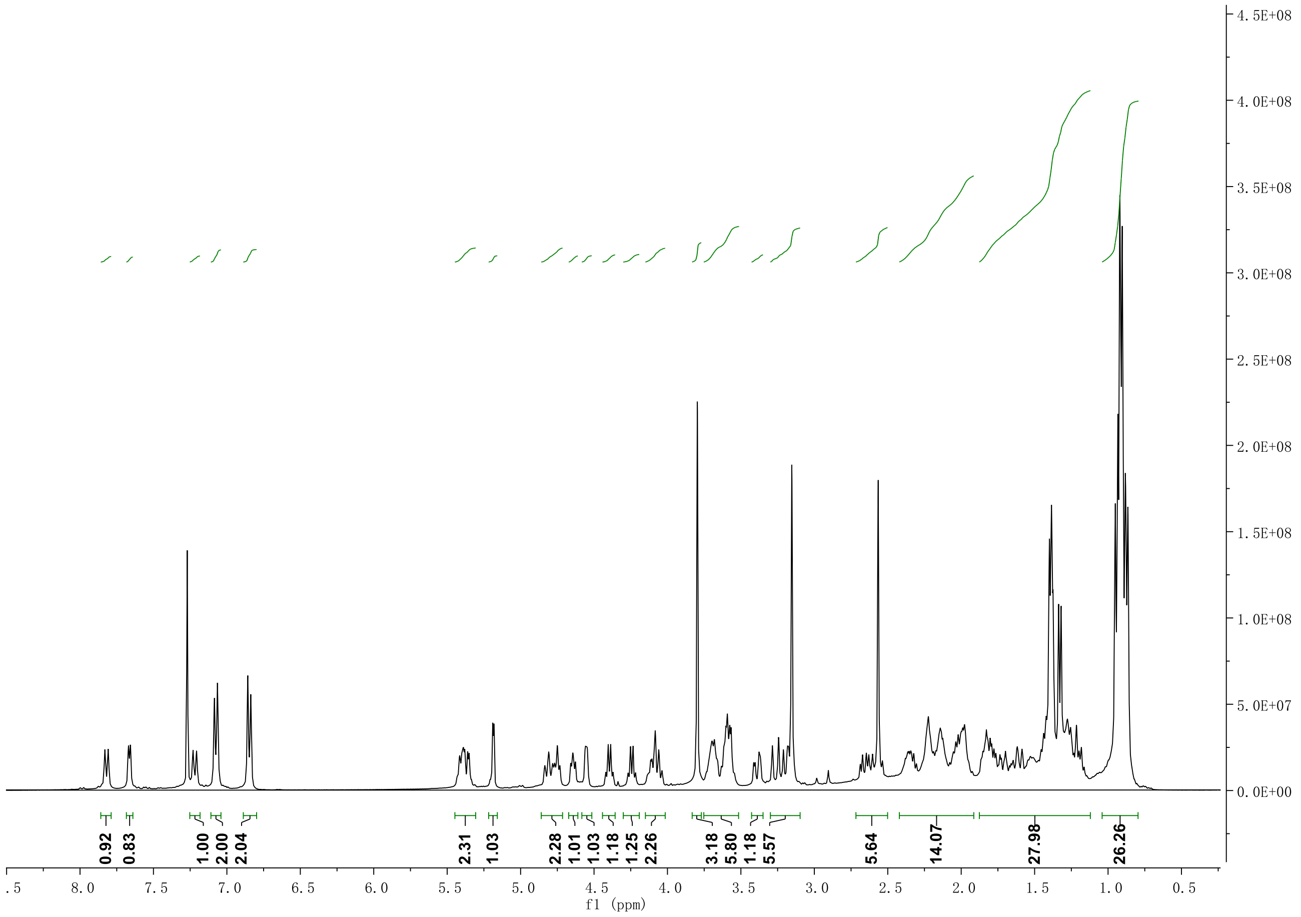

**Figure S12** $^{13}C$ NMR spectrum of didemnin B (**2**, 100 MHz). Solvent: $CDCl_3$.

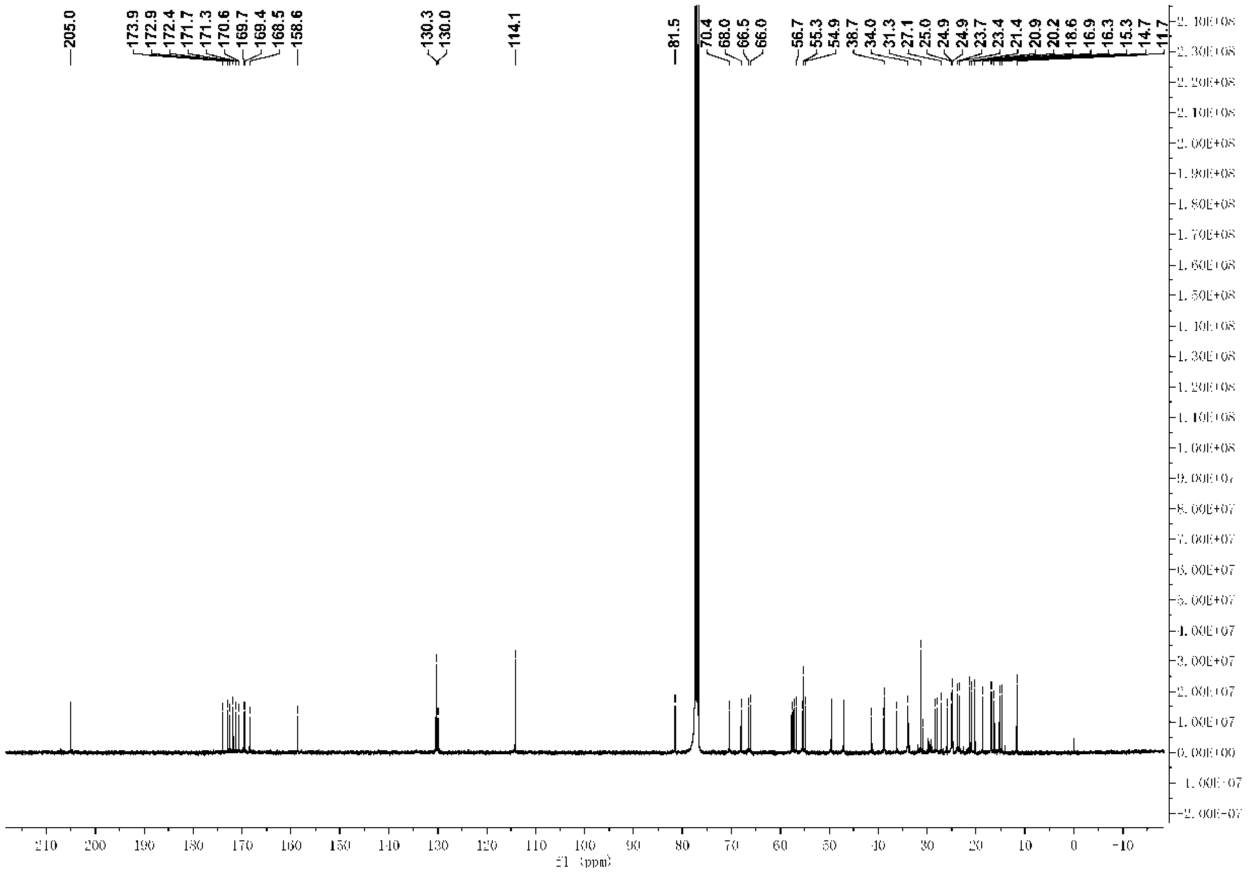

**Figure S13** $^1H$-$^1H$ COSY of didemnin B (**2**). Solvent: $CDCl_3$.

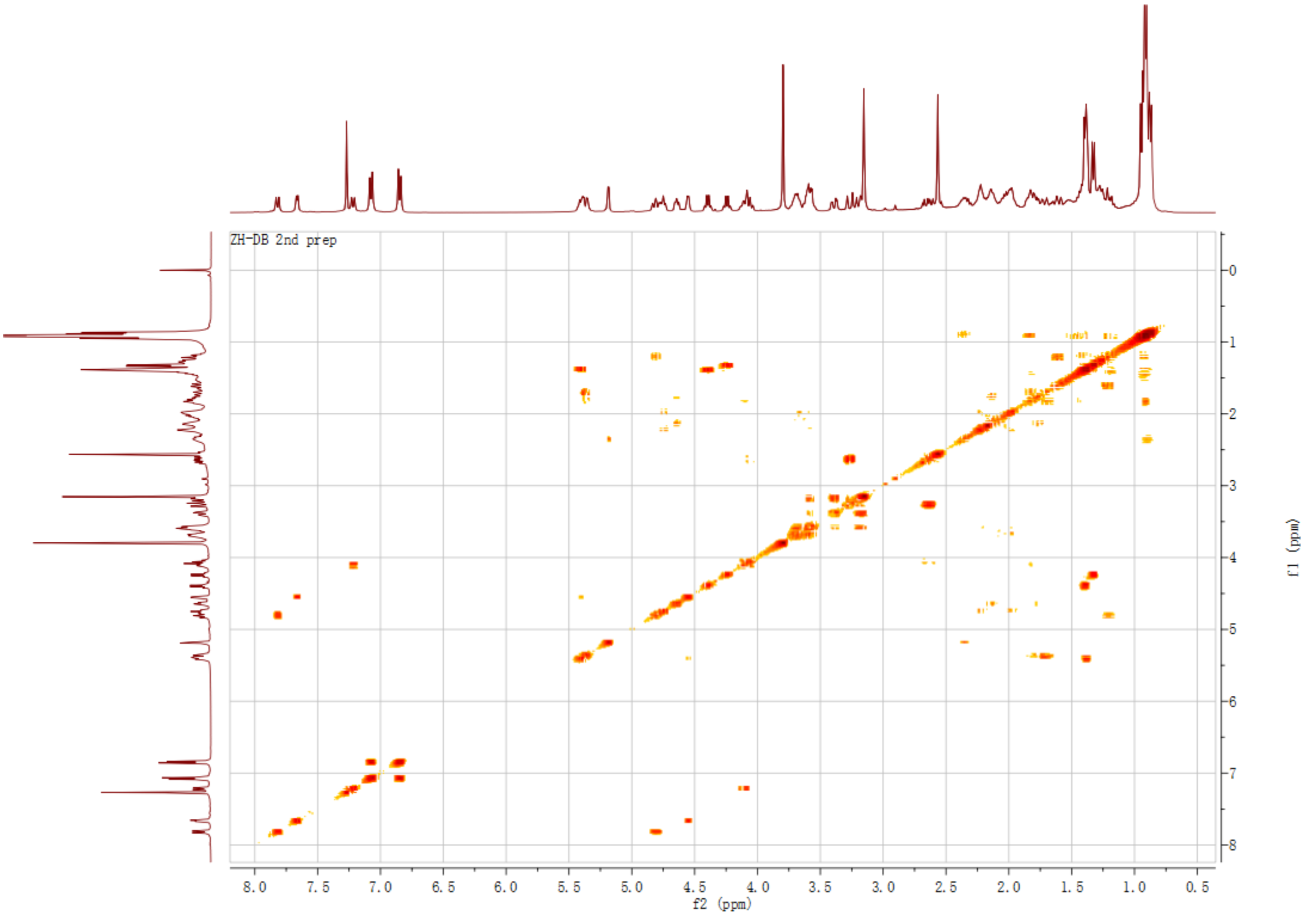

**Figure S14** $^{1}H$-$^{13}C$-HMBC NMR spectrum of didemnin B (**2**). Solvent: $CDCl_3$.

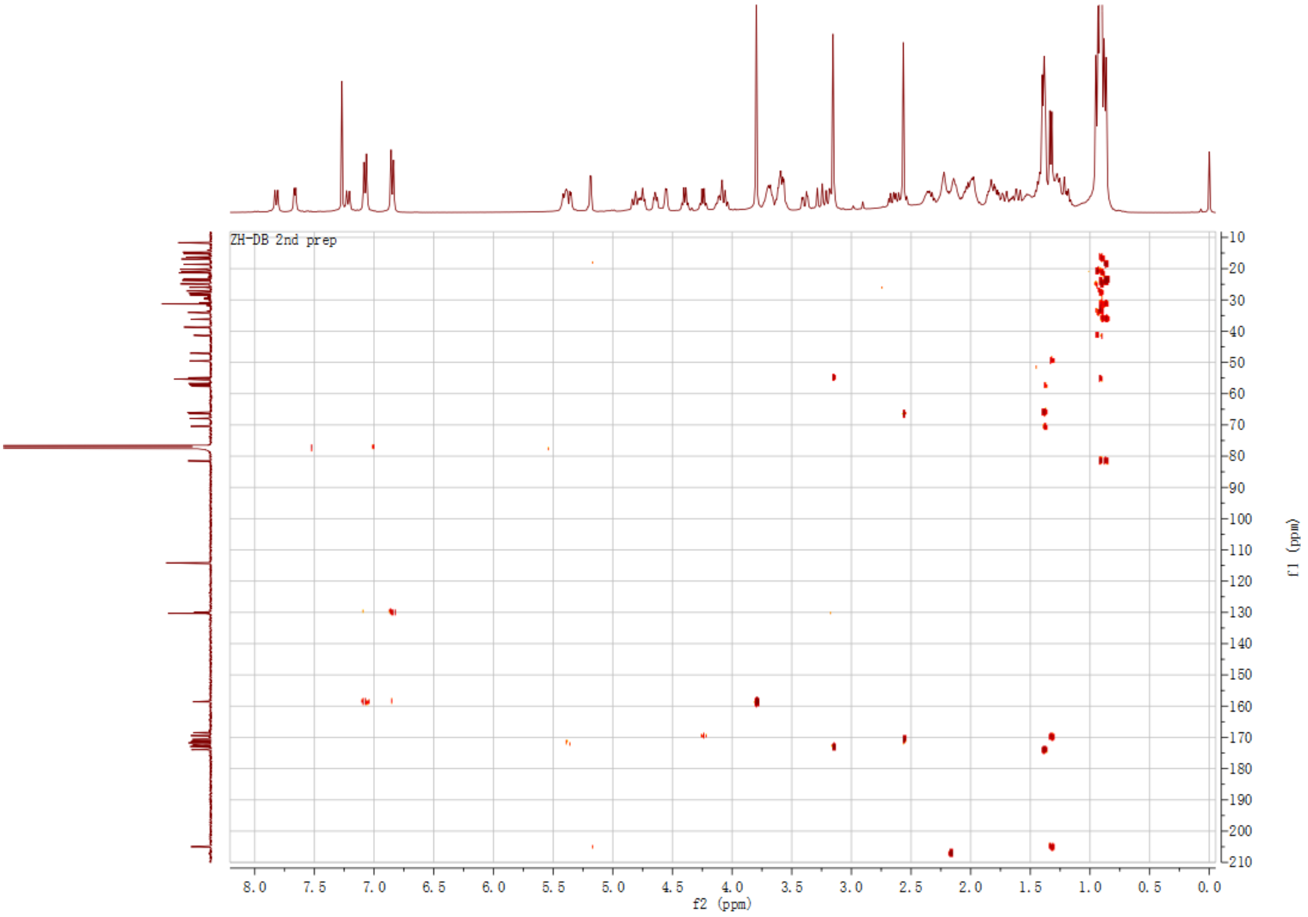

**Figure S15** $^1$H NMR spectrum of plitidepsin (**1**, 600 MHz). Solvent: $CDCl_3$.

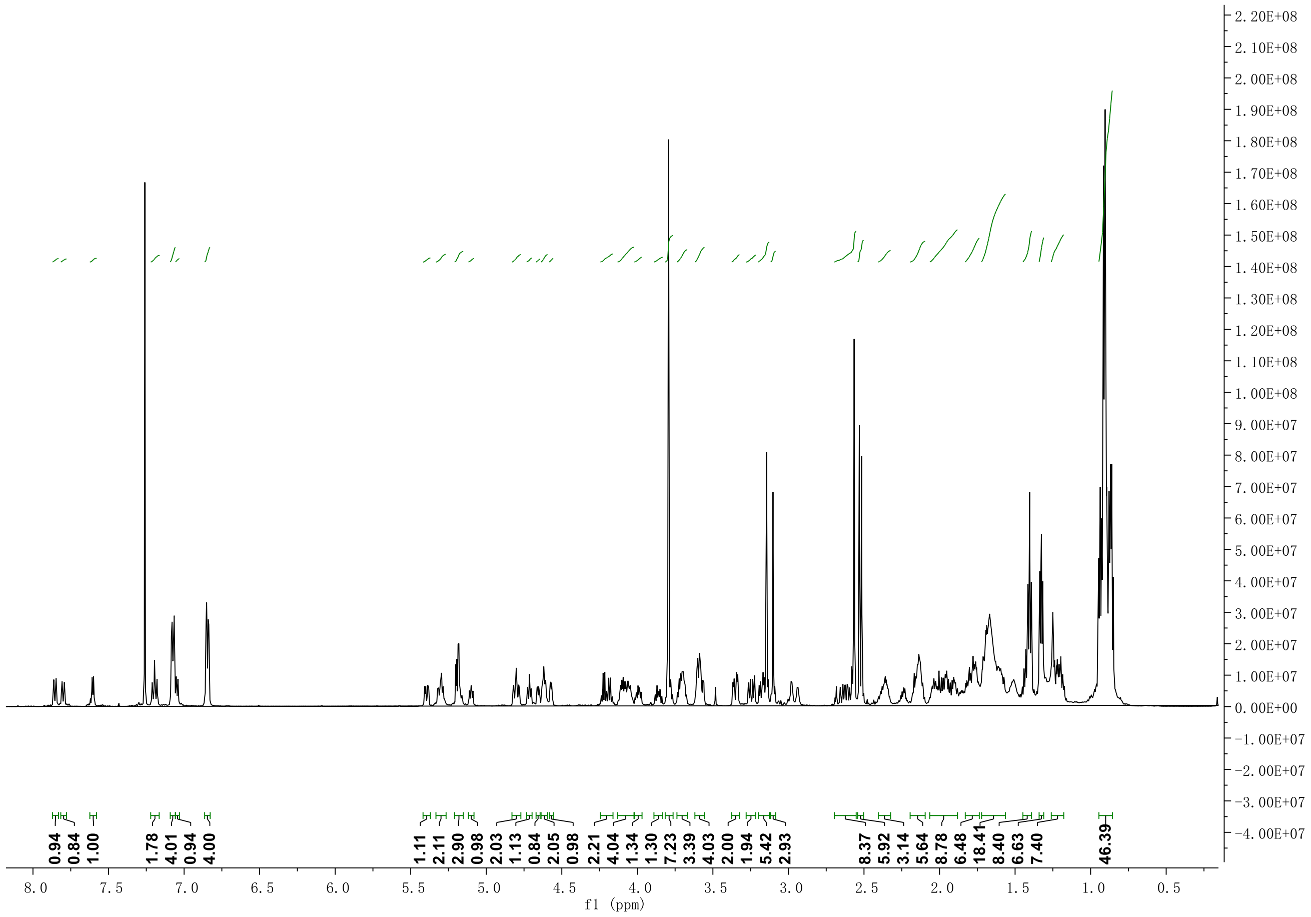

**Figure S16** $^{13}$C NMR spectrum of plitidepsin (**1**, 125 MHz). Solvent: $CDCl_3$.

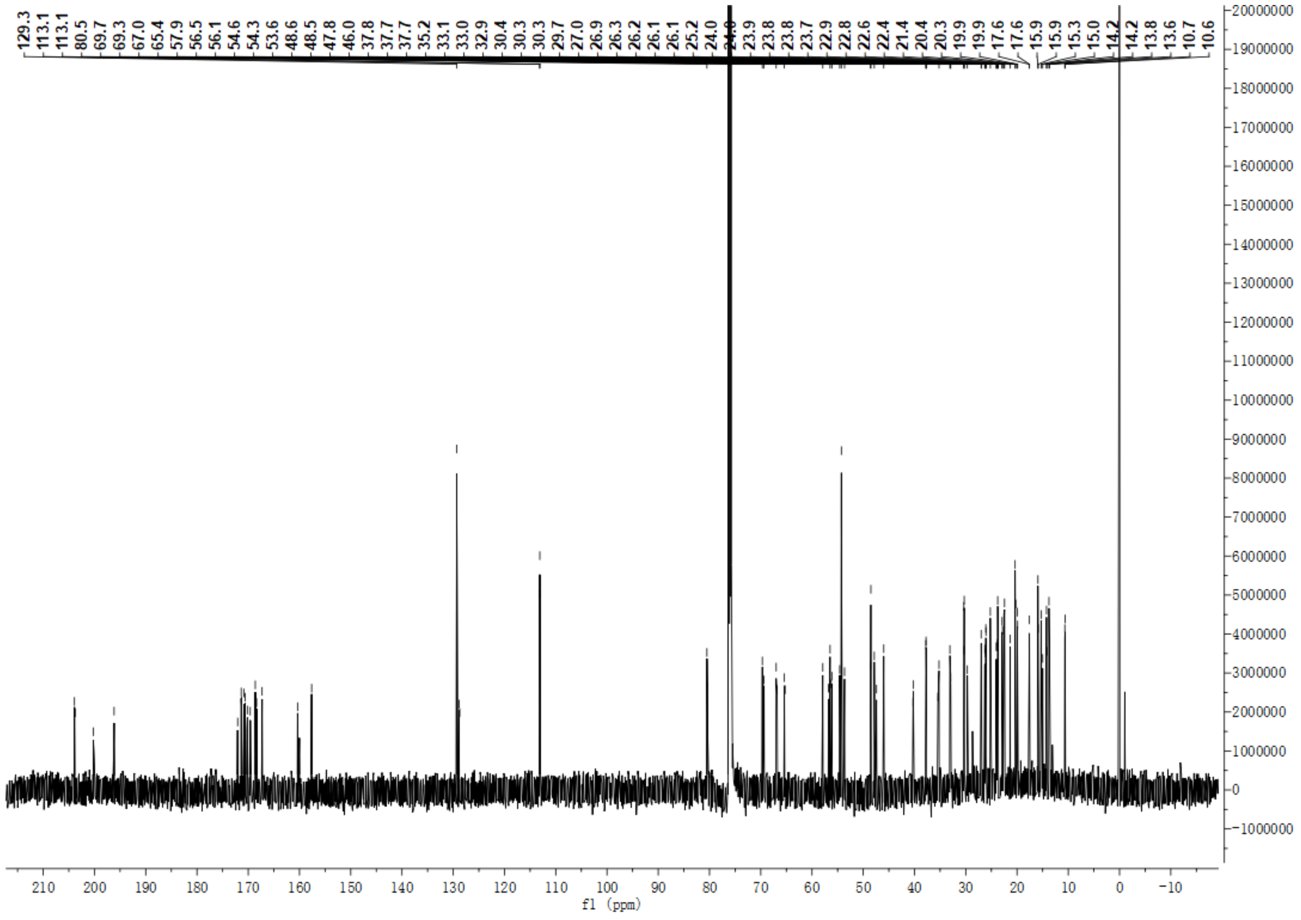

**Figure S17** $^{1}$H-$^{13}$C-HMBC of plitidepsin (**1**). Solvent: $CDCl_3$.

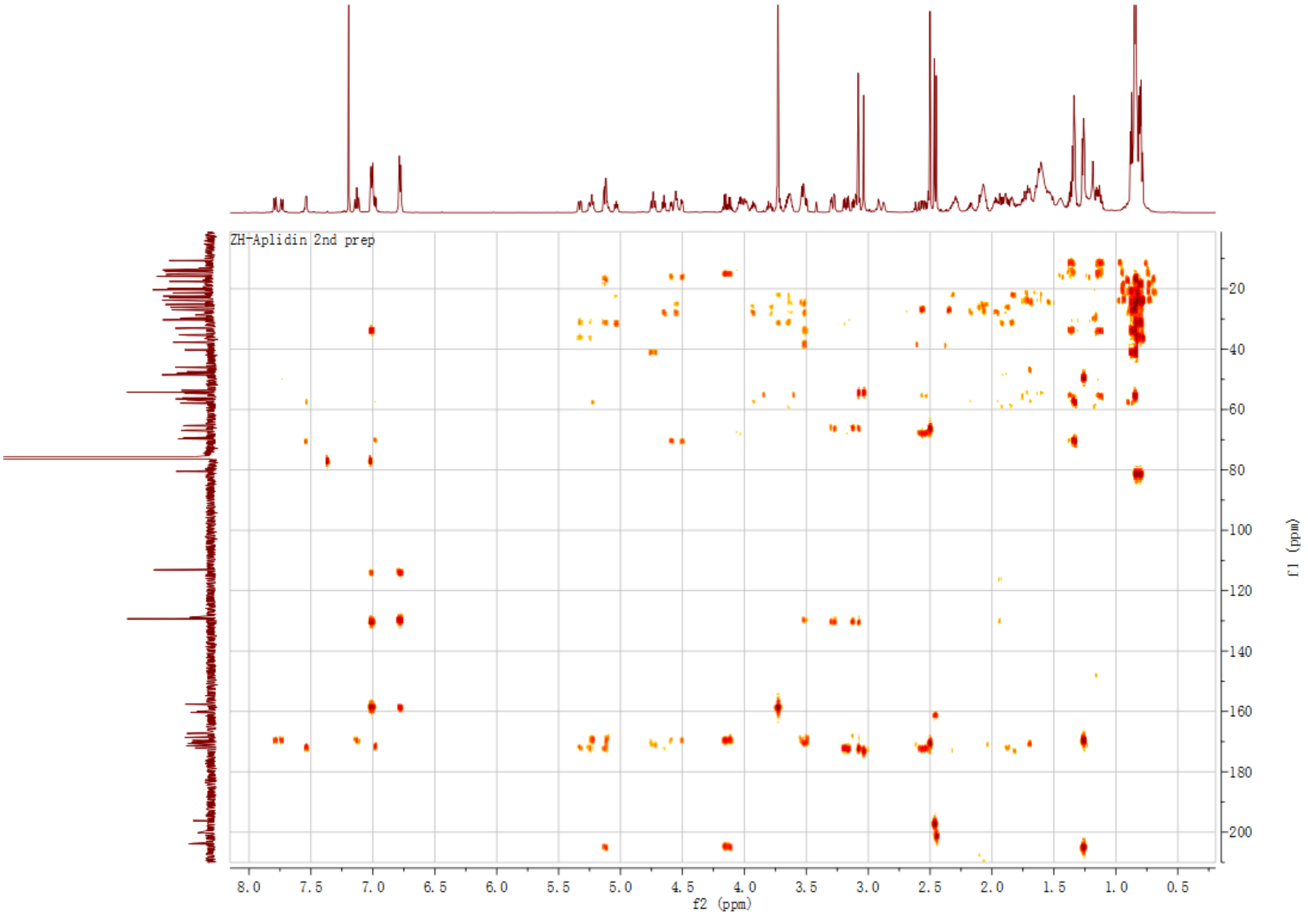

**Figure S18** $^{1}H$-$^{1}H$ COSY of plitidepsin (**1**). Solvent: $CDCl_3$.

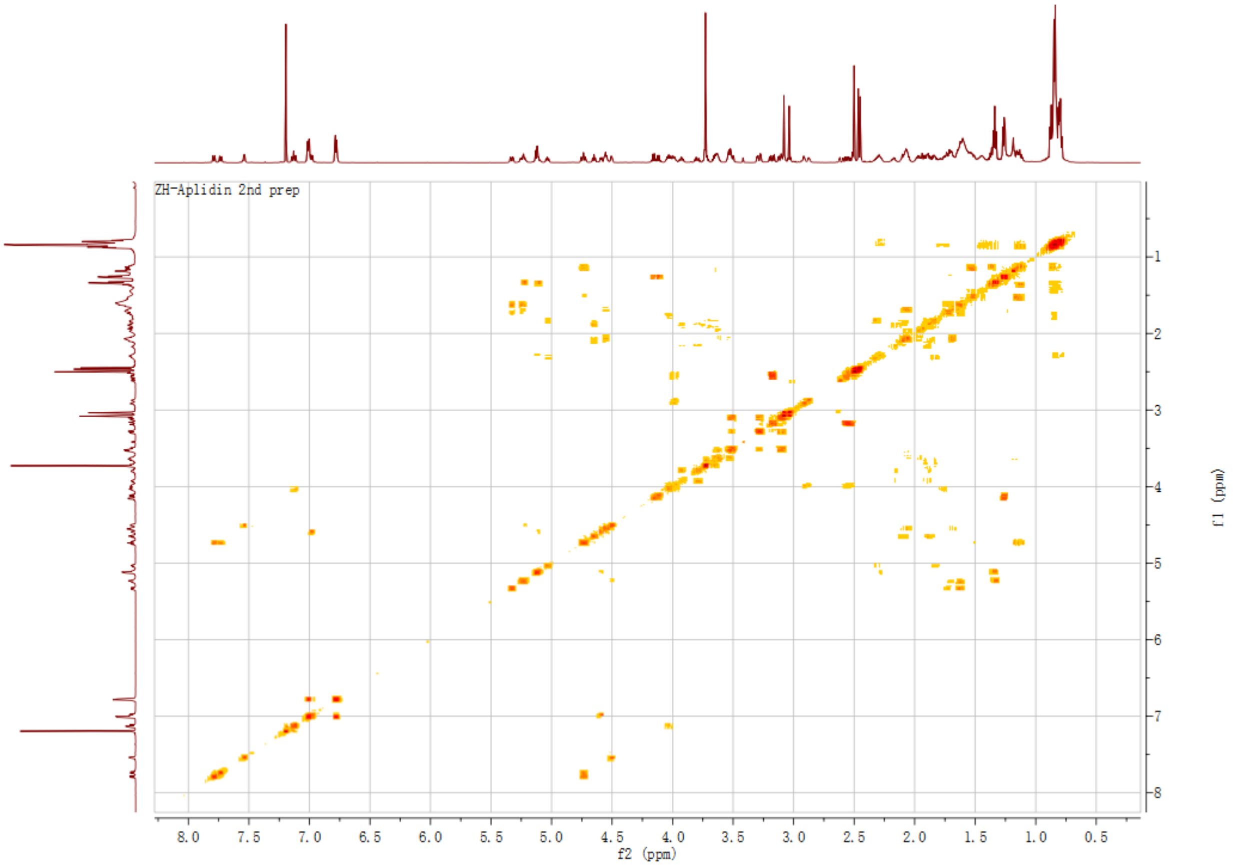

**Figure S19** $^{1}H$-$^{13}C$ HSQC of plitidepsin (**1**). Solvent: $CDCl_3$.

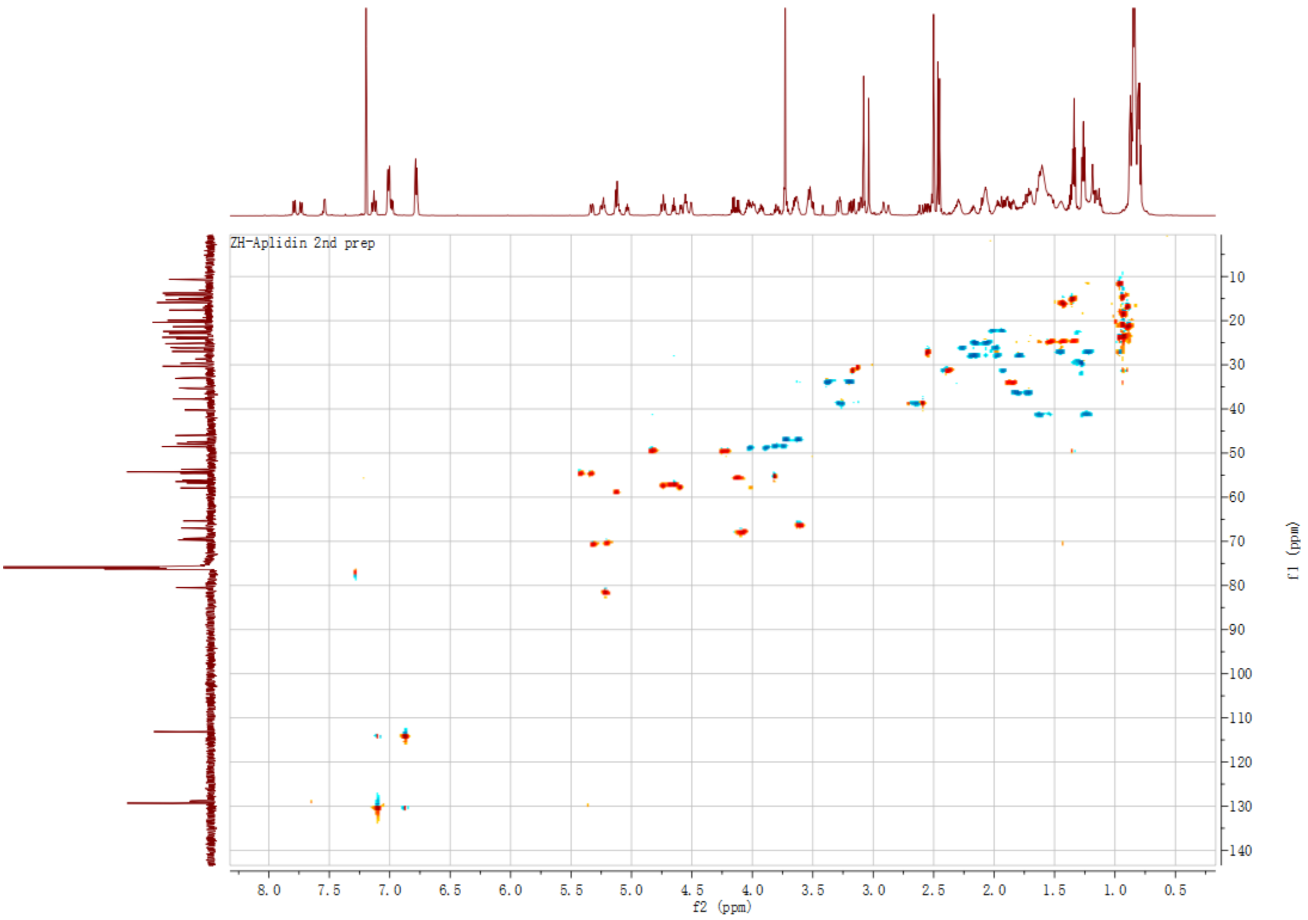

**Figure S20** $^1$H NMR of compound **5** (600 MHz). Solvent: $CDCl_3$.

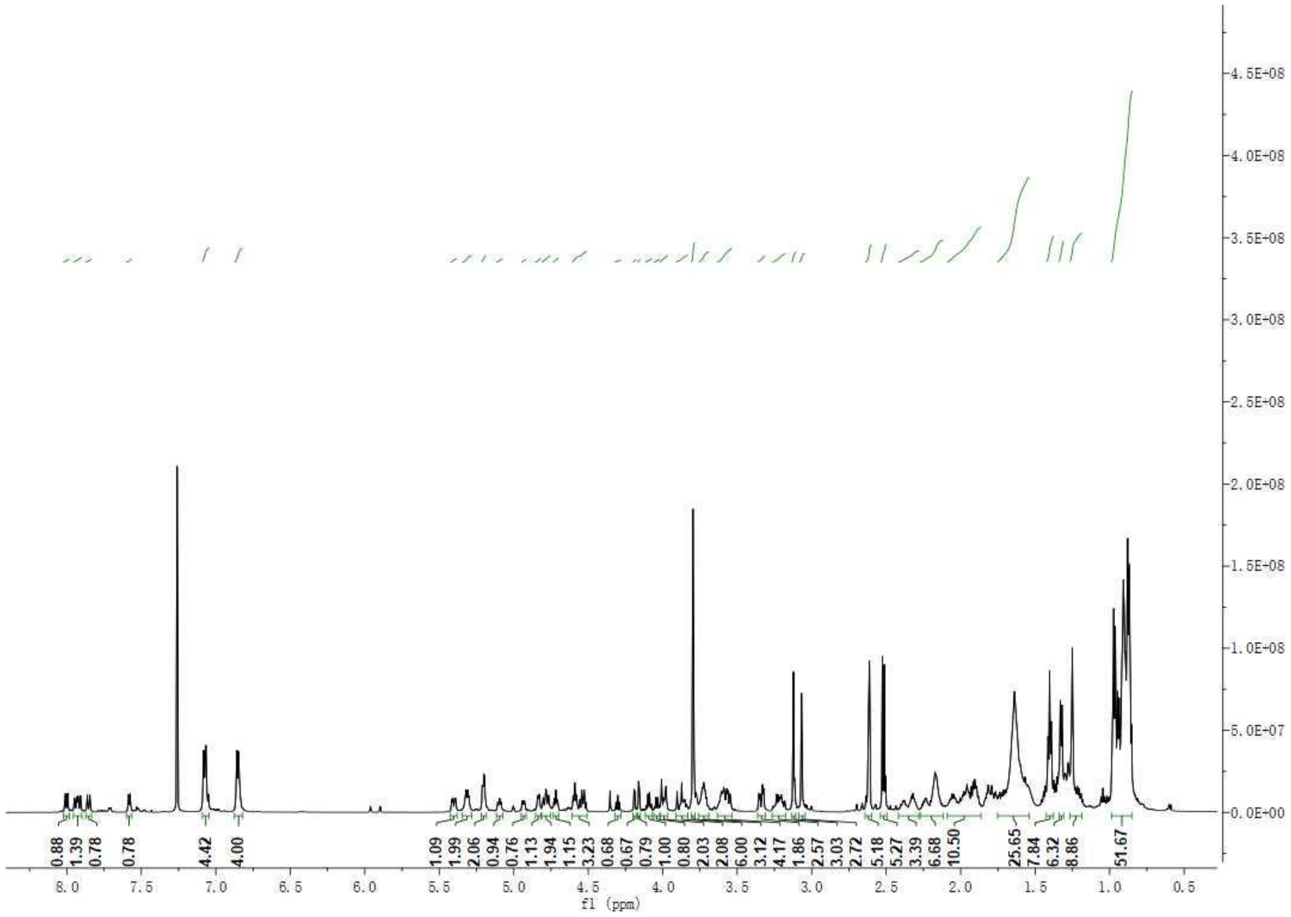

**Figure S21** $^{13}$C NMR of compound **5** (125 MHz). Solvent: $CDCl_3$.

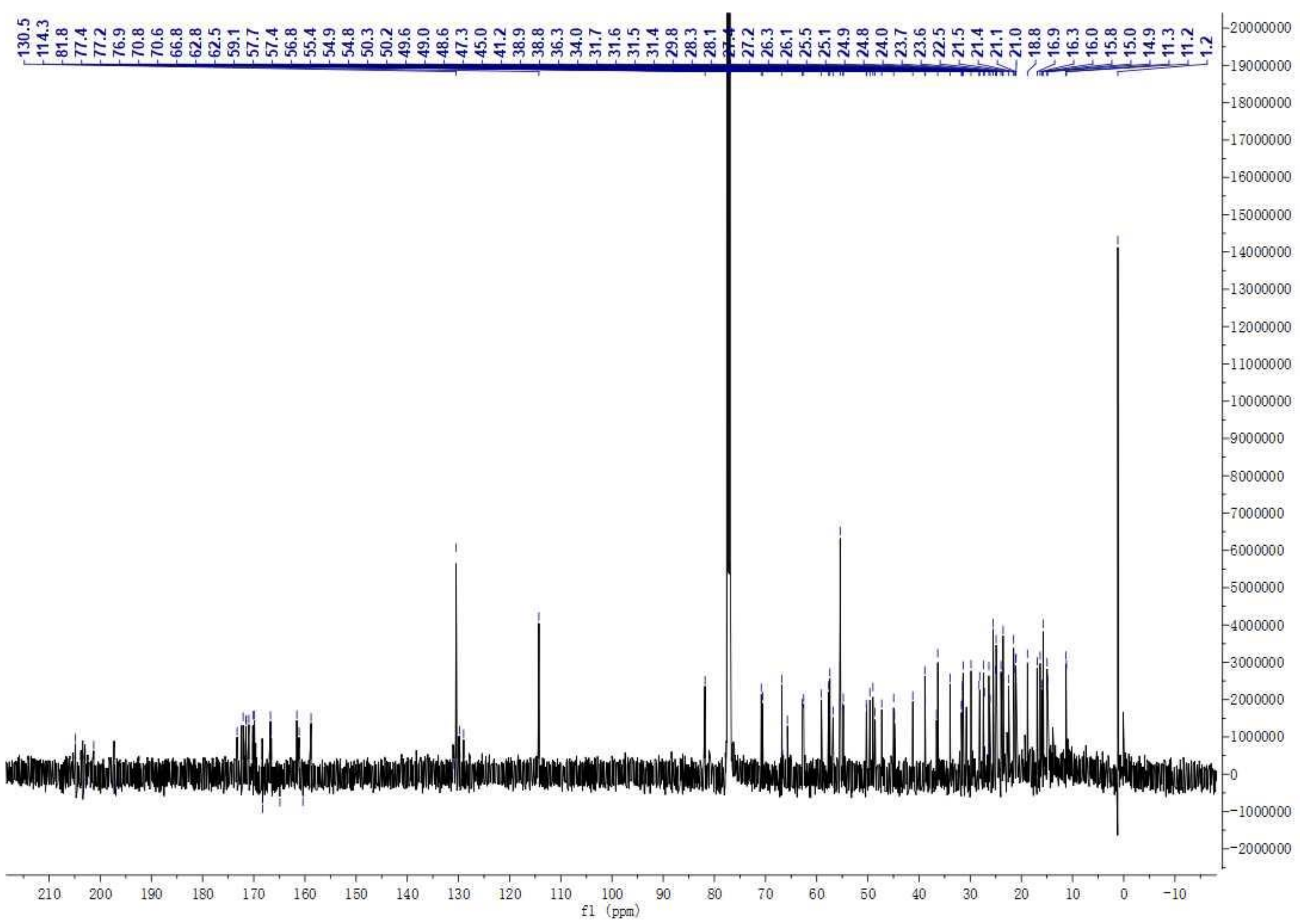

**Figure S22** $^1H$-$^1H$ COSY of compound **5**. Solvent: $CDCl_3$.

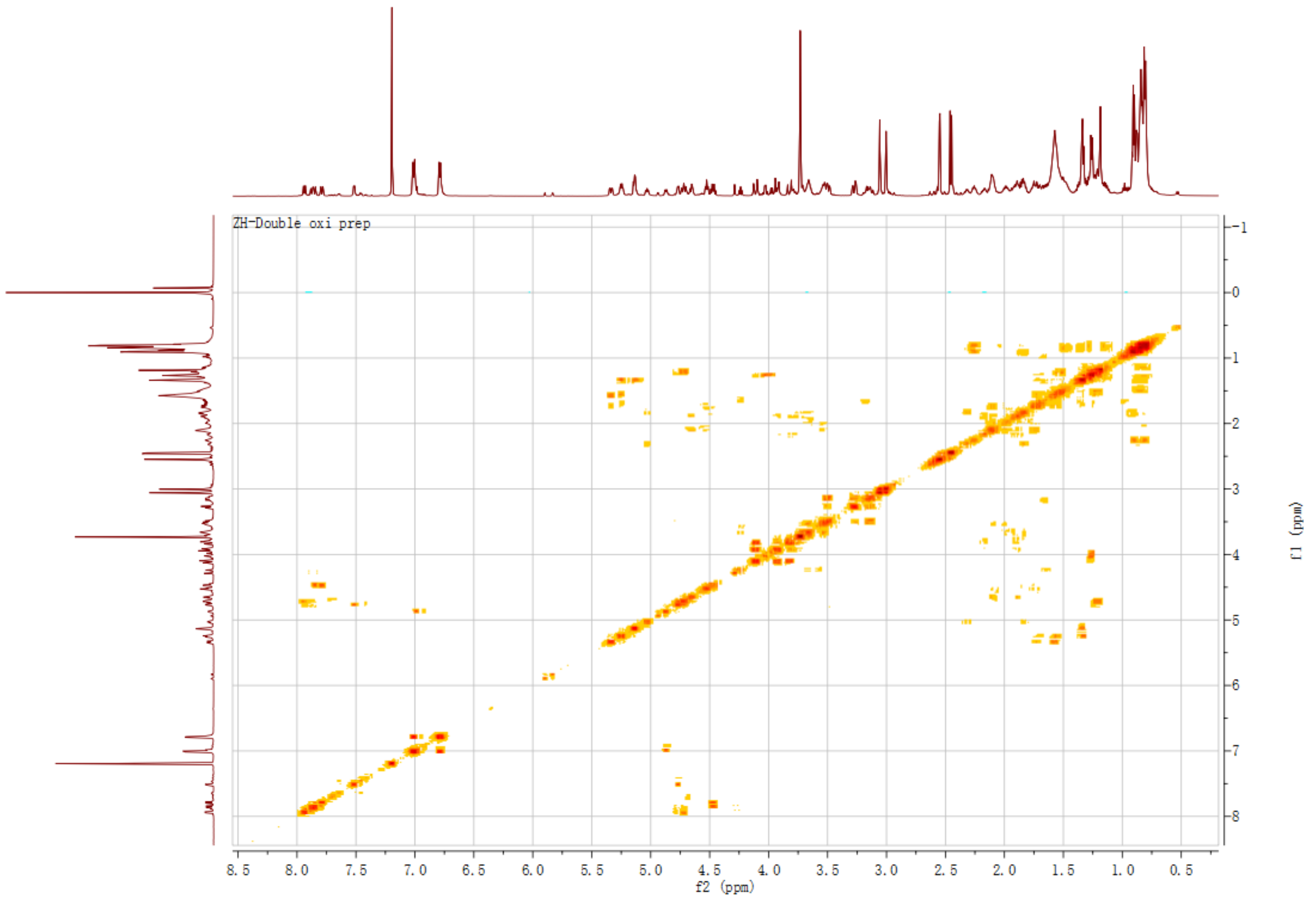

**Figure S23** $^{1}H$-$^{13}C$ HMBC of compound **5**. Solvent: $CDCl_3$.

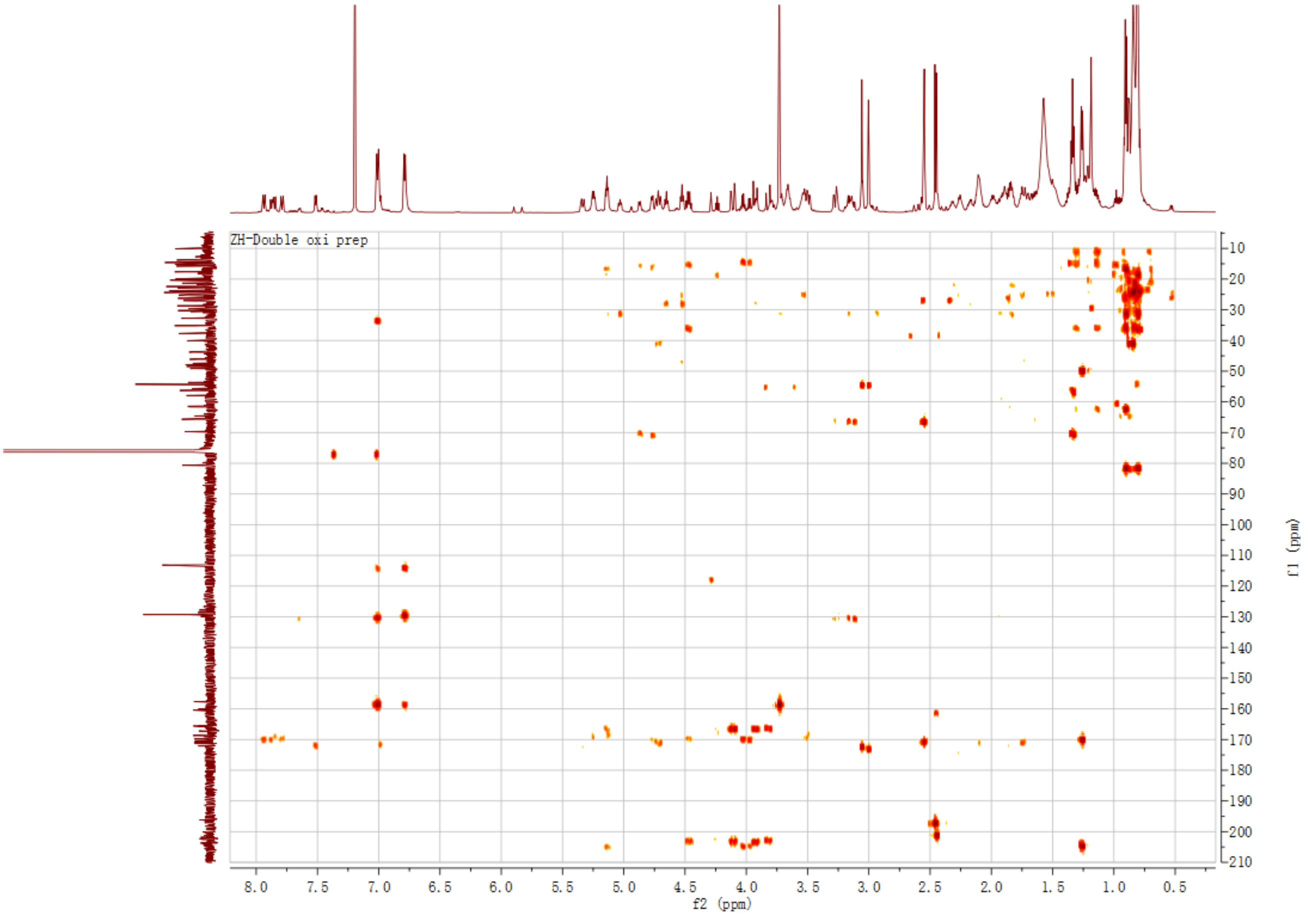

**Figure S24** $^{1}$H NMR of compound **7** (600 MHz). Solvent: $CDCl_3$.

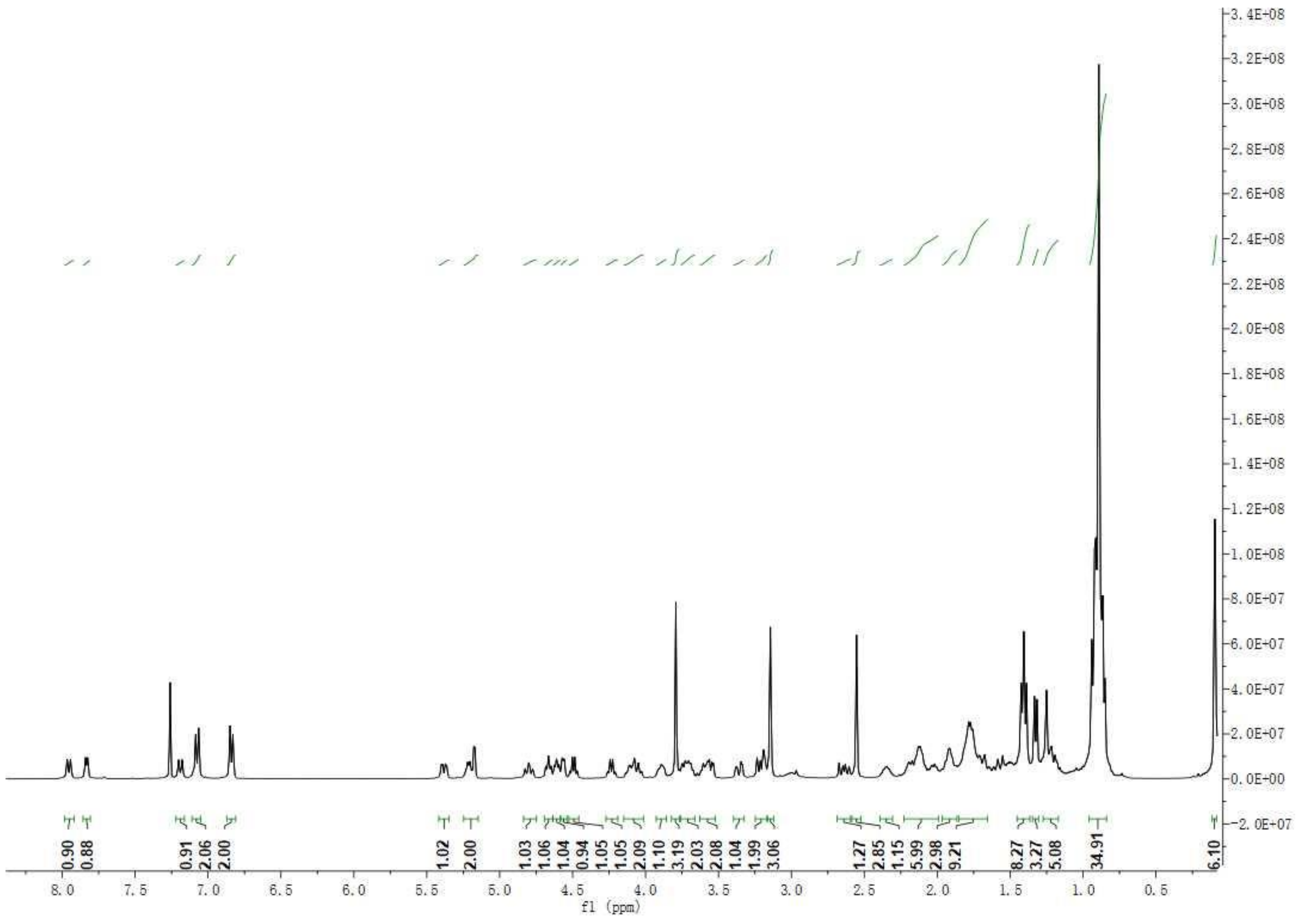

**Figure S25** $^{13}C$ NMR of compound **7** (125 MHz). Solvent: $CDCl_3$.

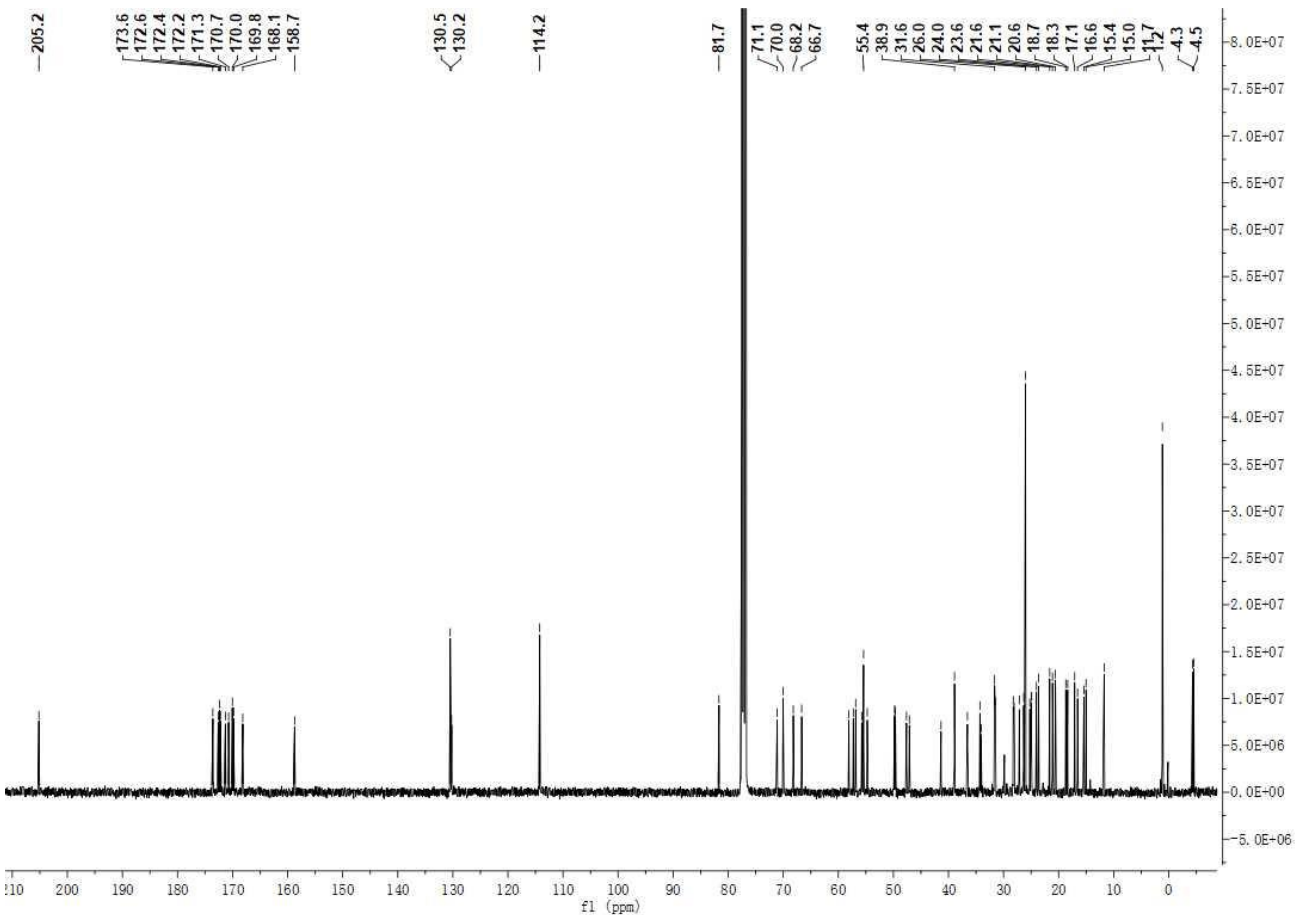

**Figure S26** $^1H$-$^1H$ COSY of compound **7** (600 MHz). Solvent: $CDCl_3$.

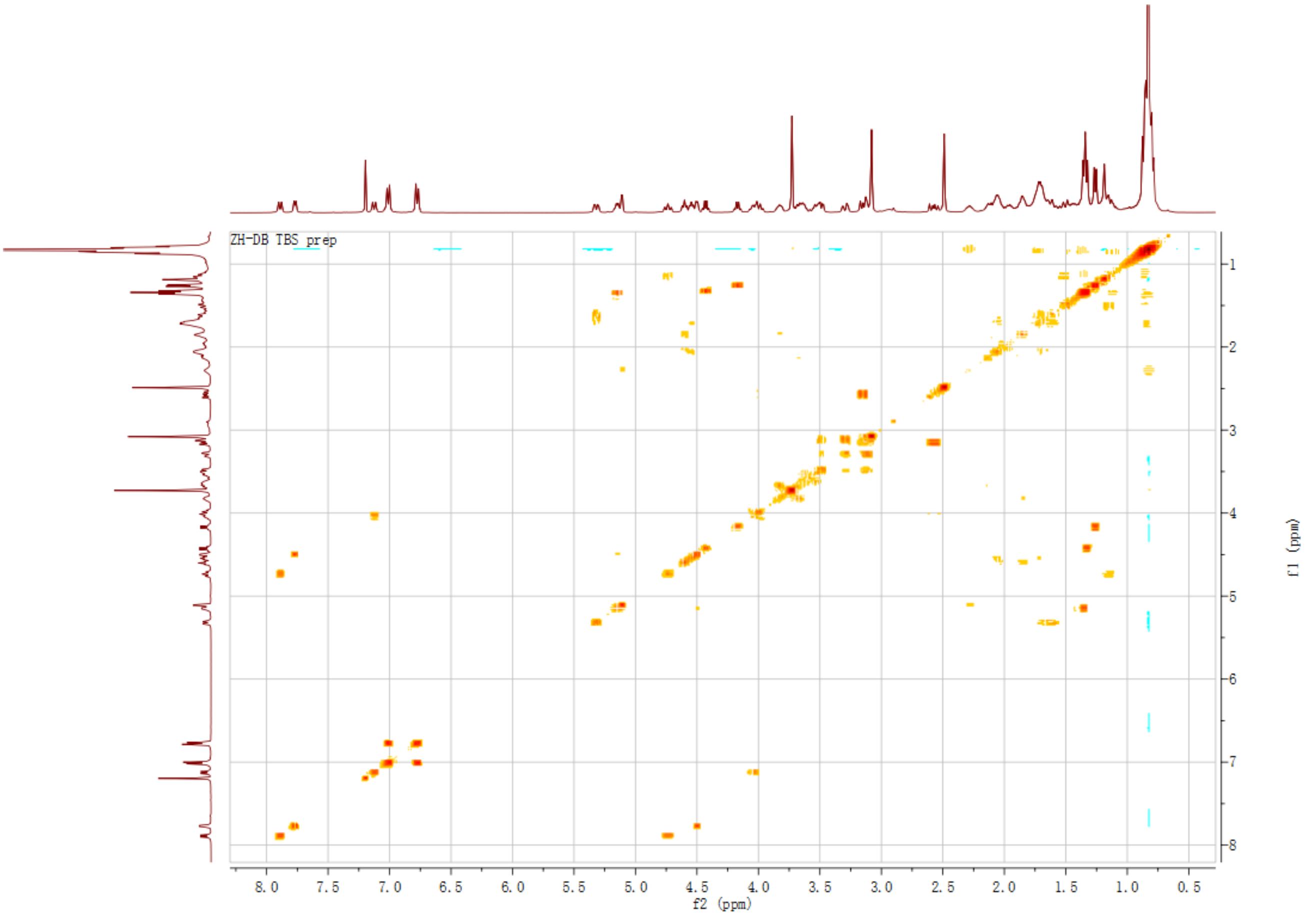

**Figure S27** $^{1}$H NMR of compound **8** (600 MHz). Solvent: $CDCl_3$.

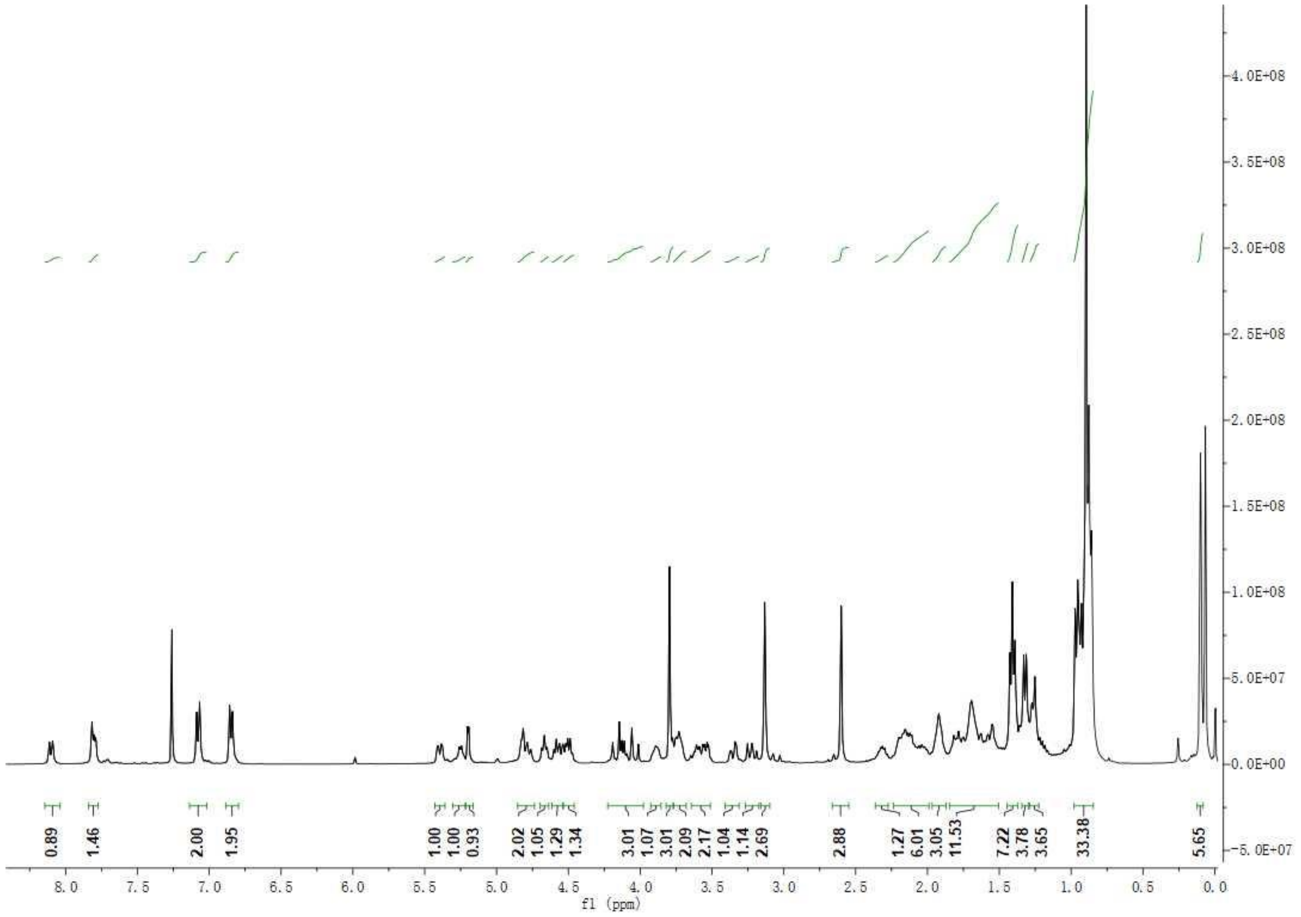

**Figure S28** $^{13}C$ NMR of compound **8** (125 MHz). Solvent: $CDCl_3$.

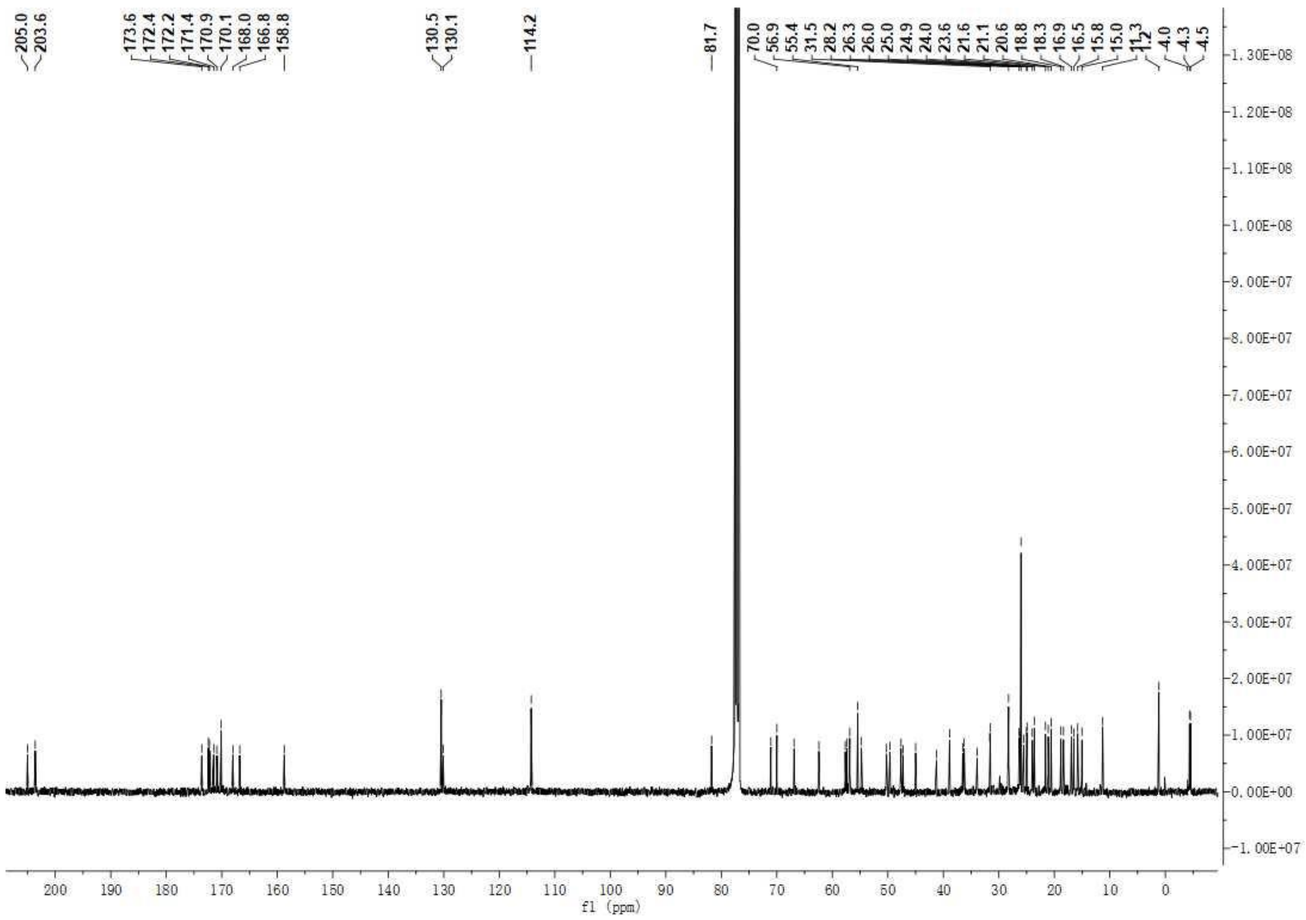

**Figure S29** $^{1}$H NMR spectrum of compound **6** (400 MHz). Solvent: $CDCl_3$.

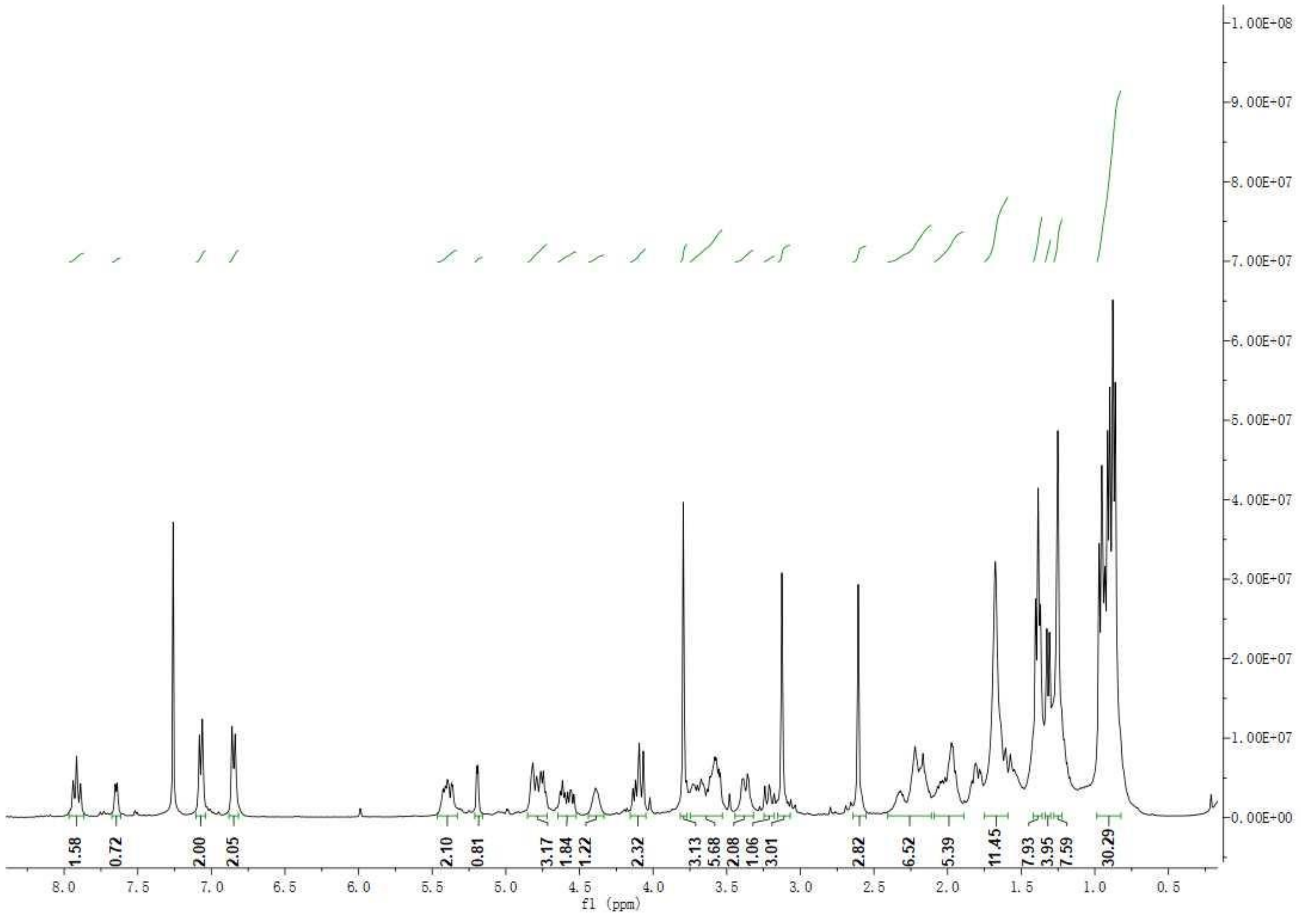

**Figure S30** $^{13}$C NMR spectrum of compound **6** (100 MHz). Solvent: $CDCl_3$.

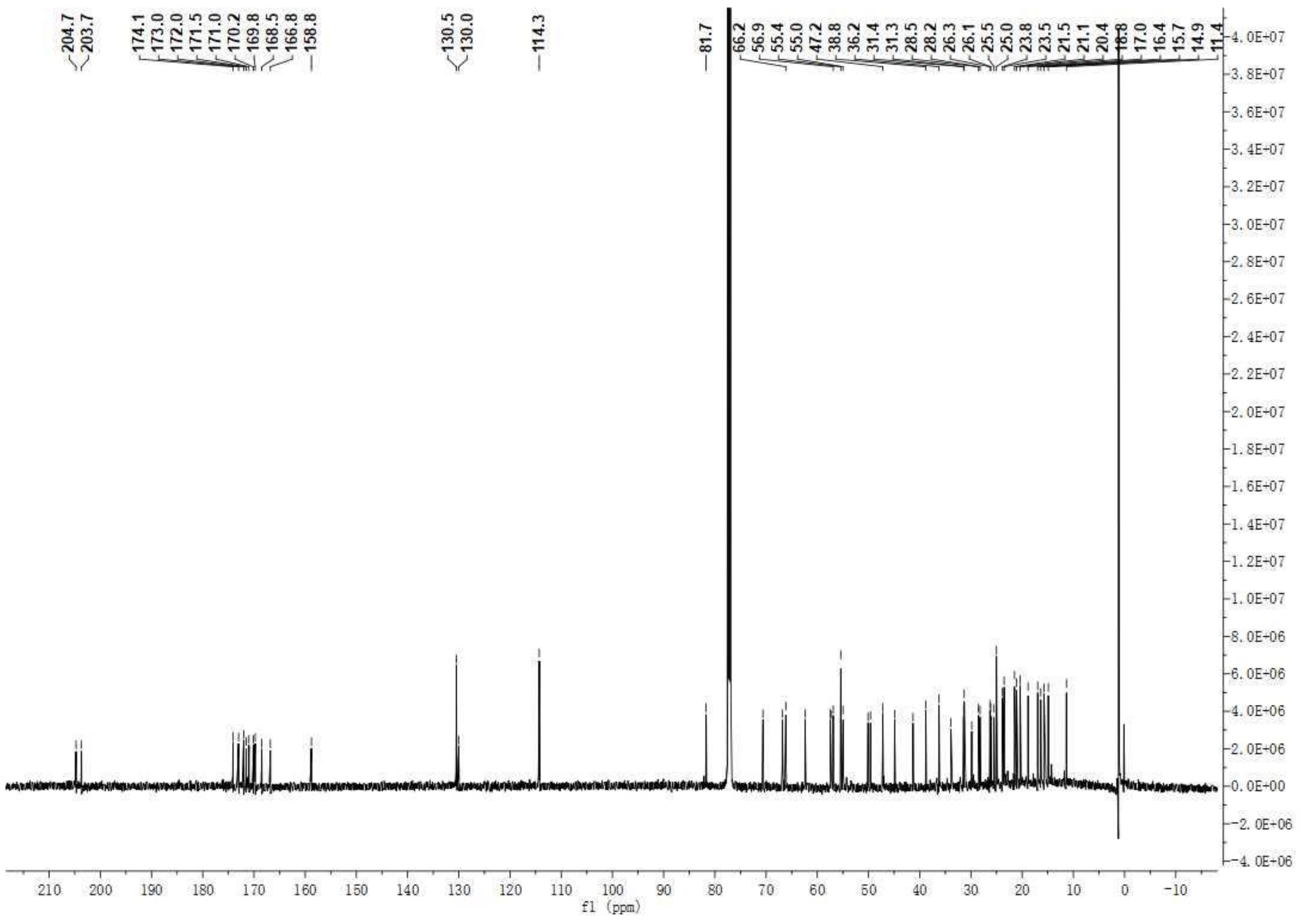

**Figure S31** $^{1}H$-$^{13}C$ HMBC spectrum of compound **6**. Solvent: $CDCl_3$.

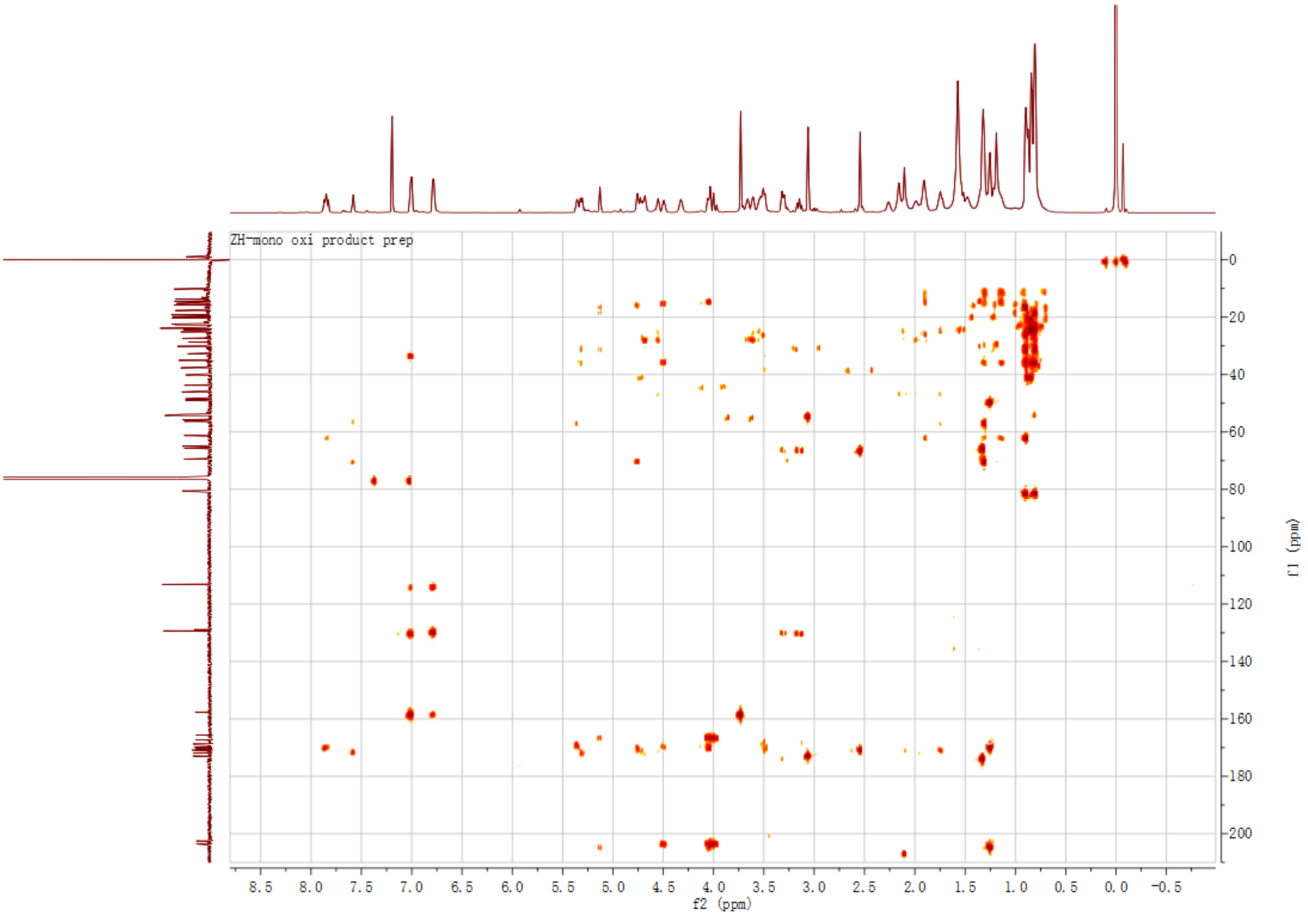

**Figure S32** $^1$H-$^1$H COSY spectrum of compound **6**. Solvent: $CDCl_3$.

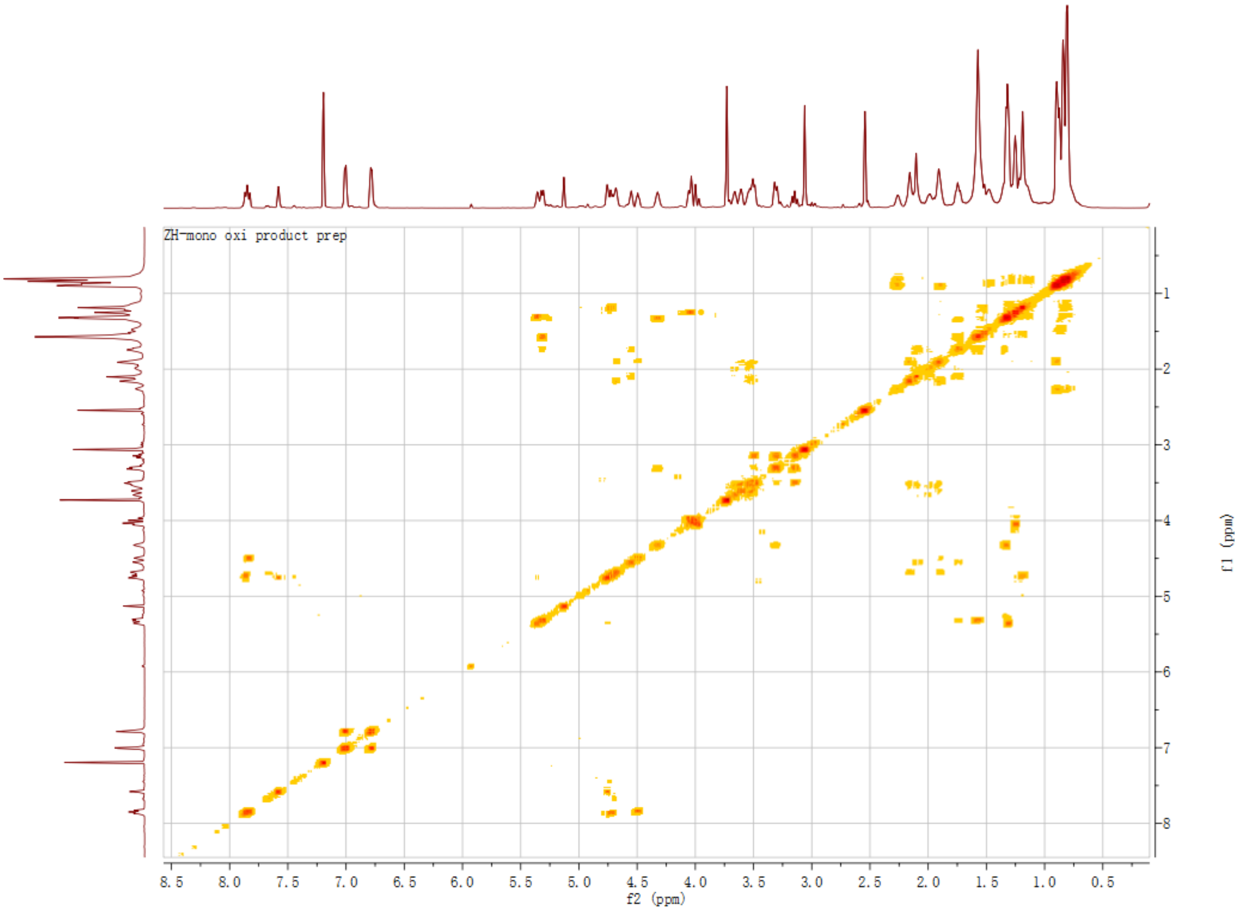

**Figure S33** $^{1}H$-$^{13}C$ HSQC spectrum of compound **6**. Solvent: $CDCl_3$.

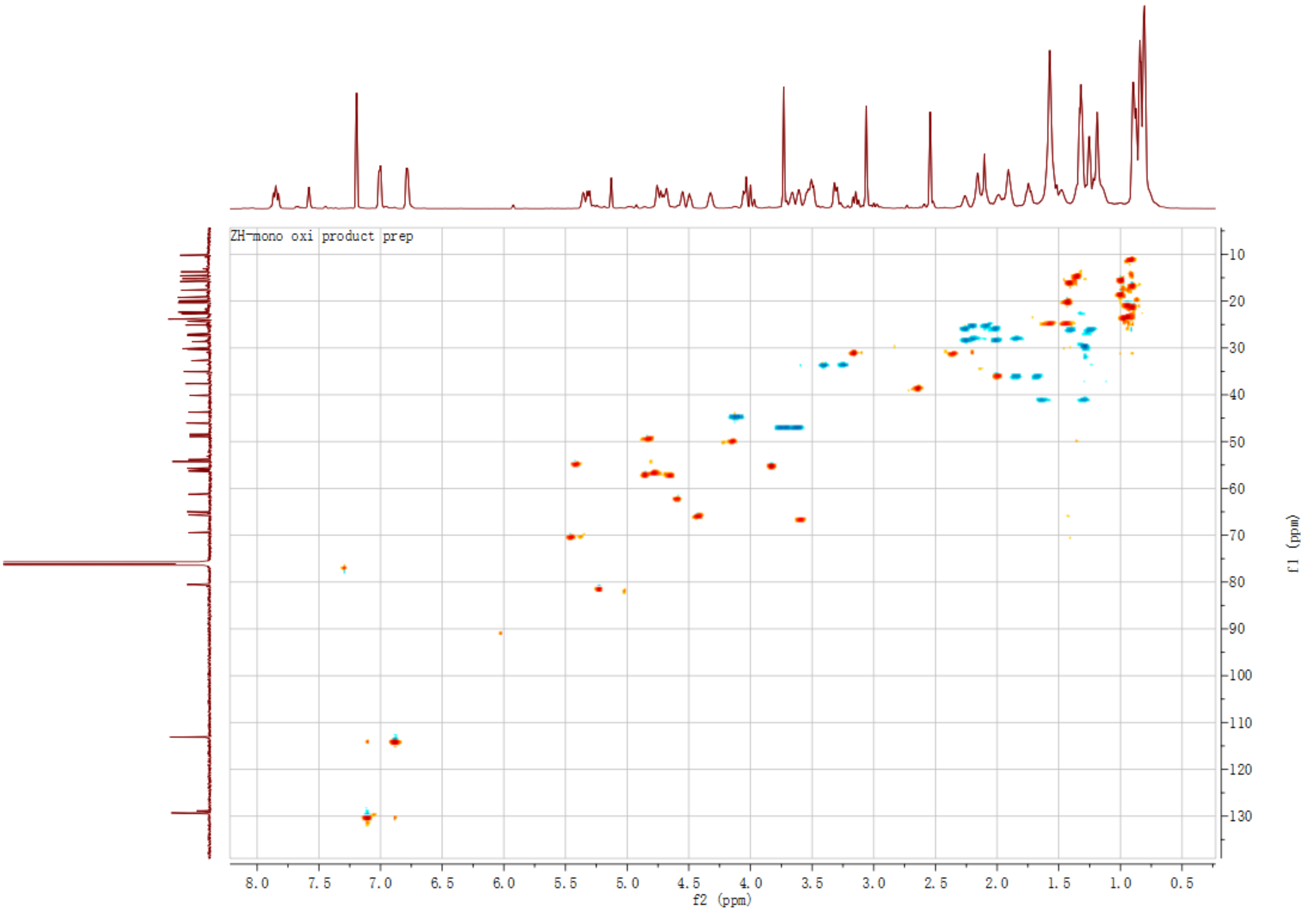

**Figure S34** HRMS and MS² analysis of didemnin B (**2**).

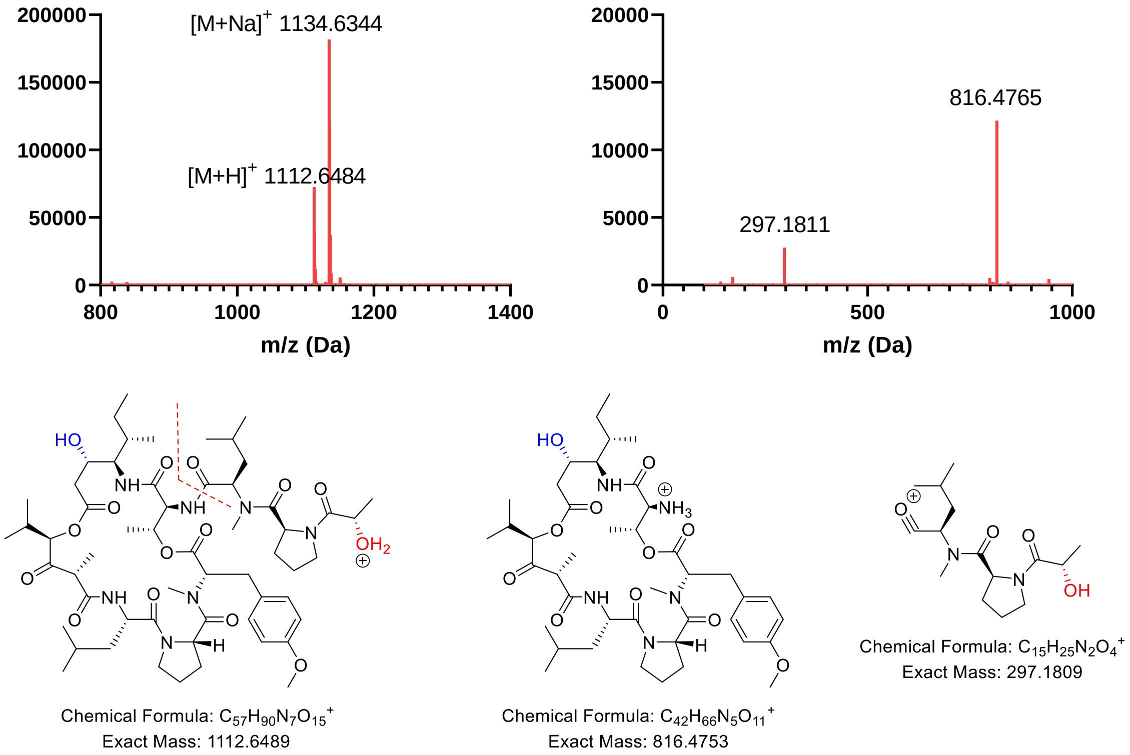


**Figure S35** MS and MS² analysis of plitidepsin (**1**).

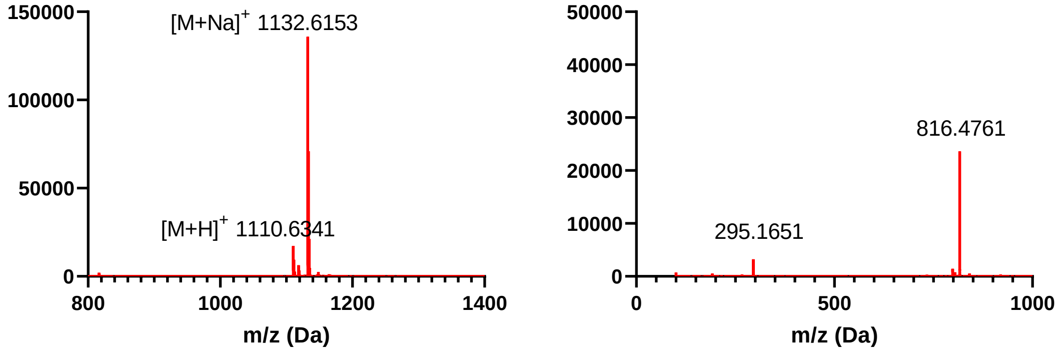


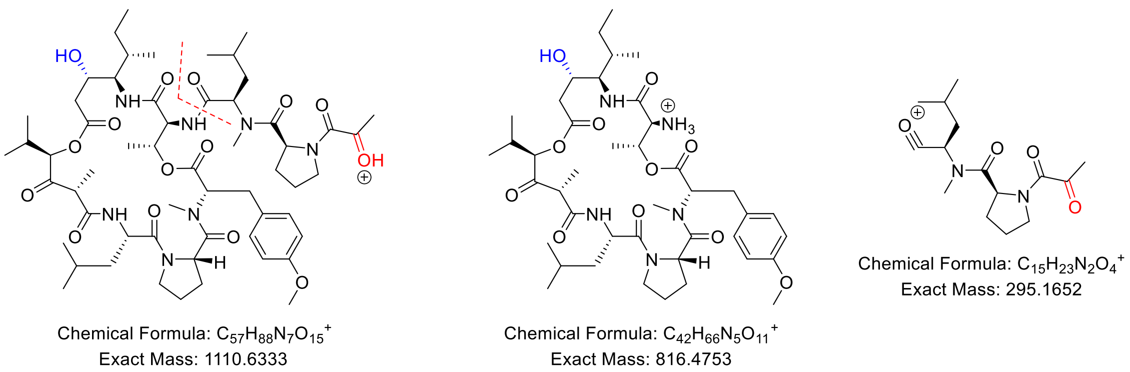


**Figure S36** MS and MS$^2$ analysis of compound **5**.

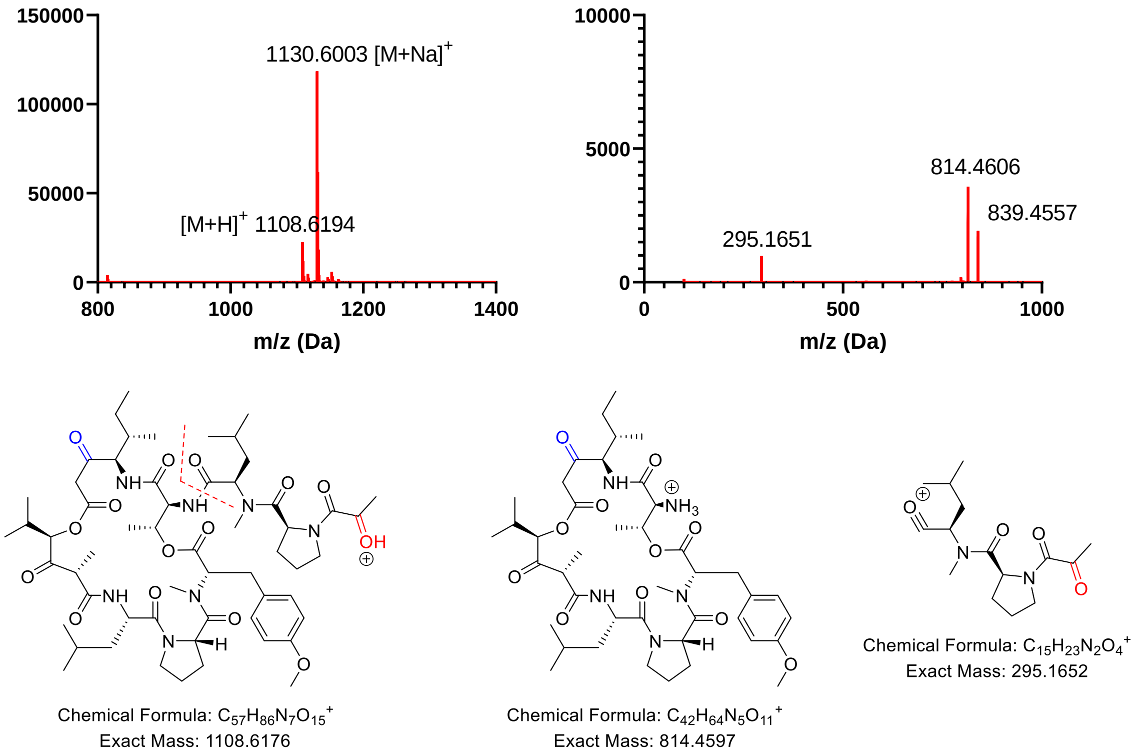


**Figure S37** MS and MS$^2$ analysis of compound **6**.

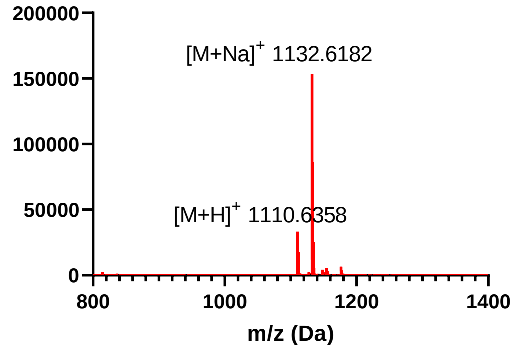


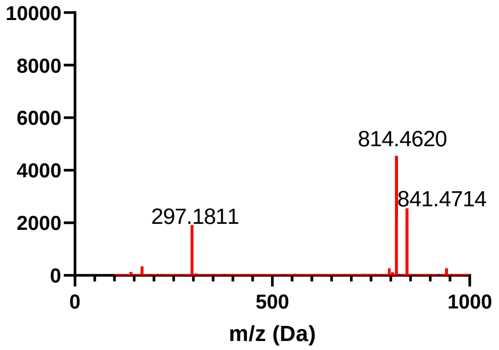


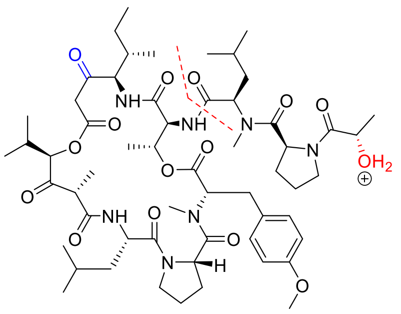

Chemical Formula: $C_{57}H_{88}N_7O_{15}^+$
Exact Mass: 1110.6333

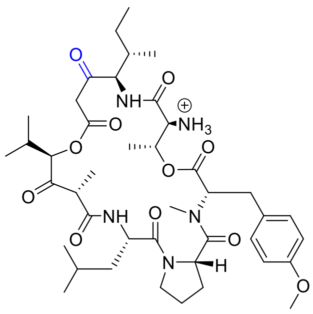

Chemical Formula: $C_{42}H_{64}N_5O_{11}^+$
Exact Mass: 814.4597

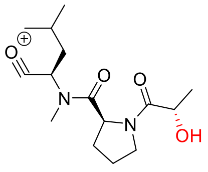

Chemical Formula: $C_{15}H_{25}N_2O_4^+$
Exact Mass: 297.1809

**Figure S38** MS and MS$^2$ analysis of compound **3**.

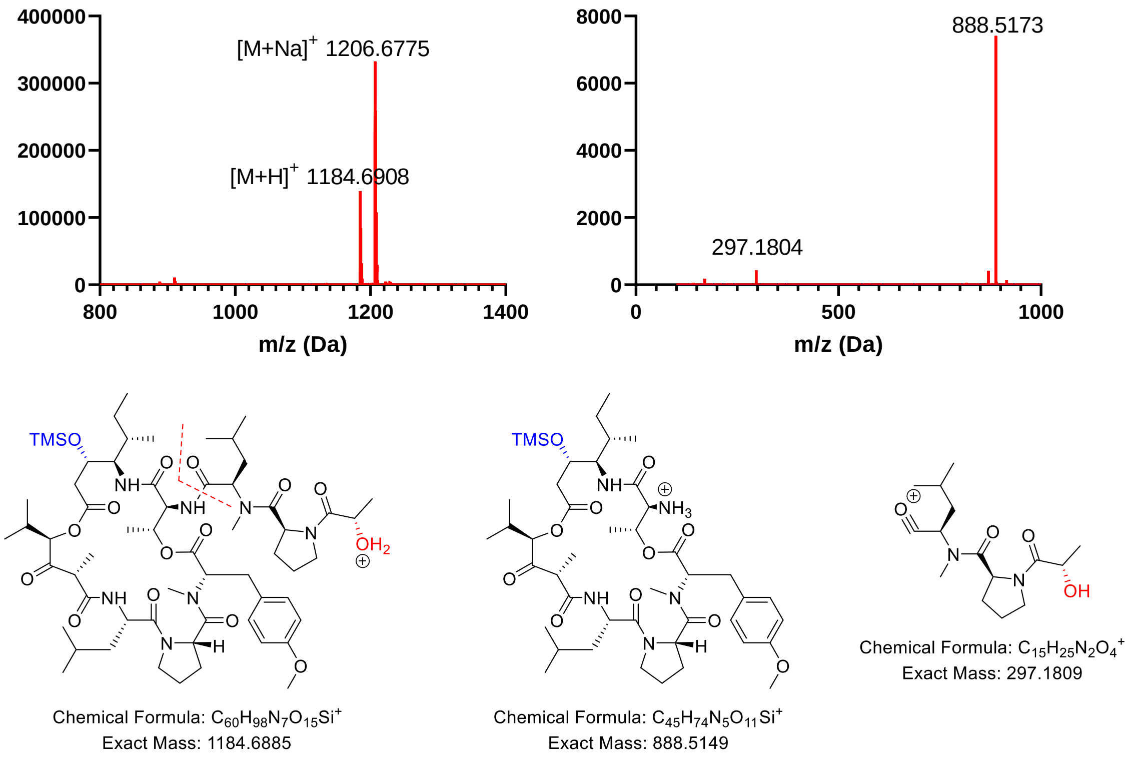


**Figure S39** MS and $MS^2$ analysis of compound **4**.

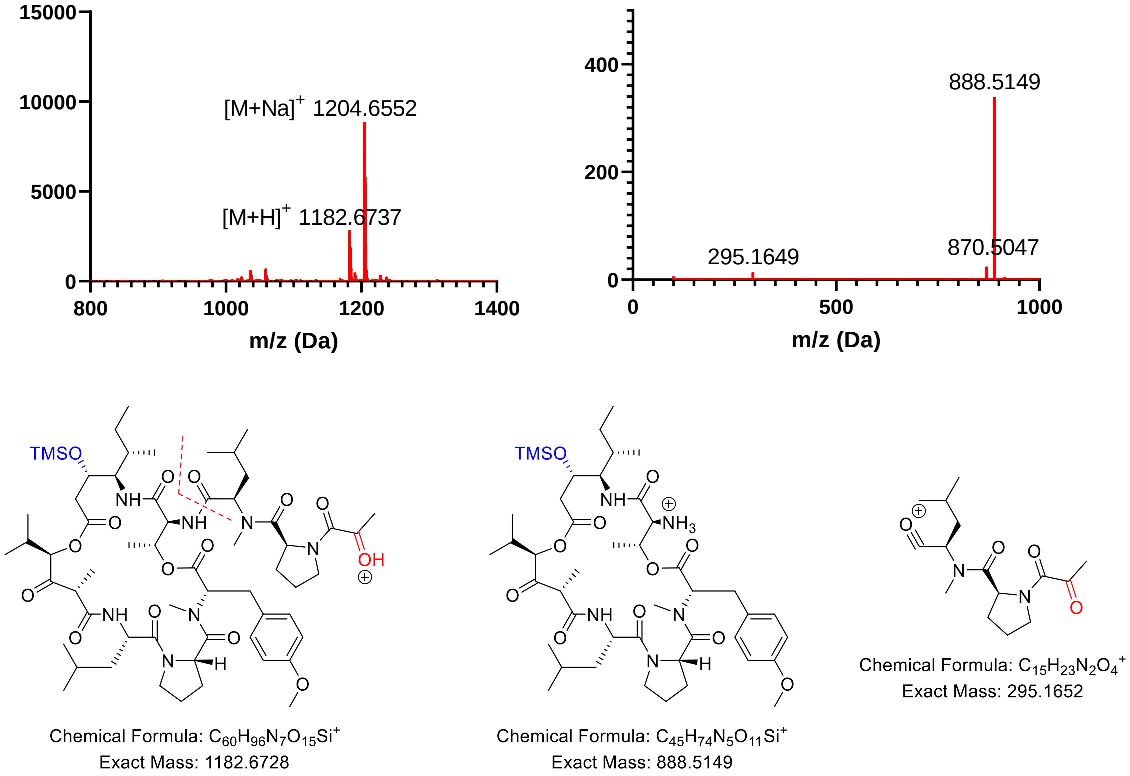


**Figure S40** MS and $MS^2$ analysis of compound **7**.

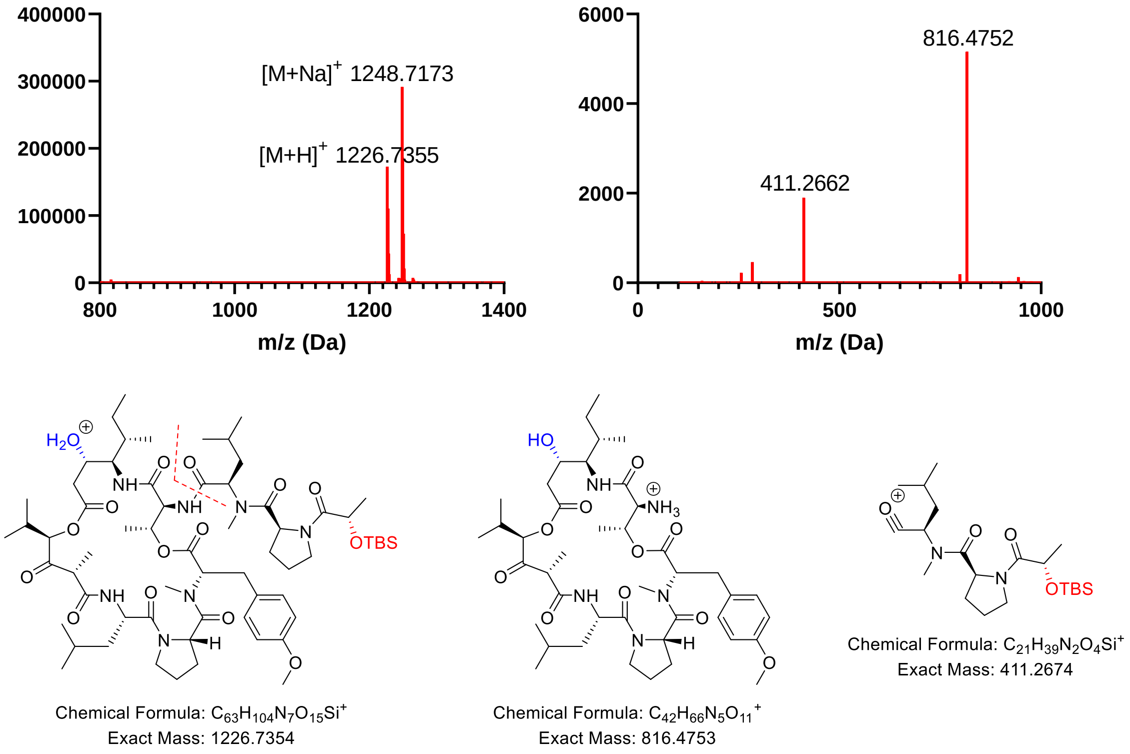


**Figure S41** MS and MS$^2$ analysis of compound **8**.

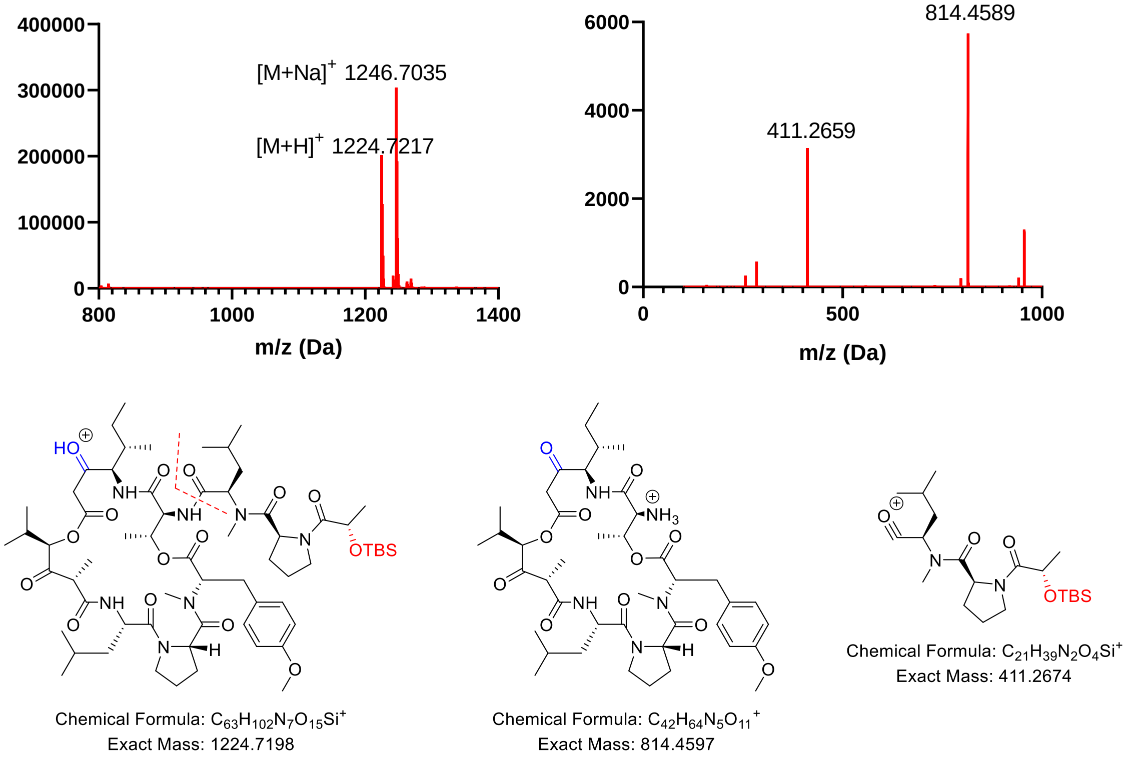


**Figure S42** LC-HRMS chromatography, LC-ELSD chromatography and UV spectra (210 nm) of the purified compounds **1**, **2**, **5**, and **6**.

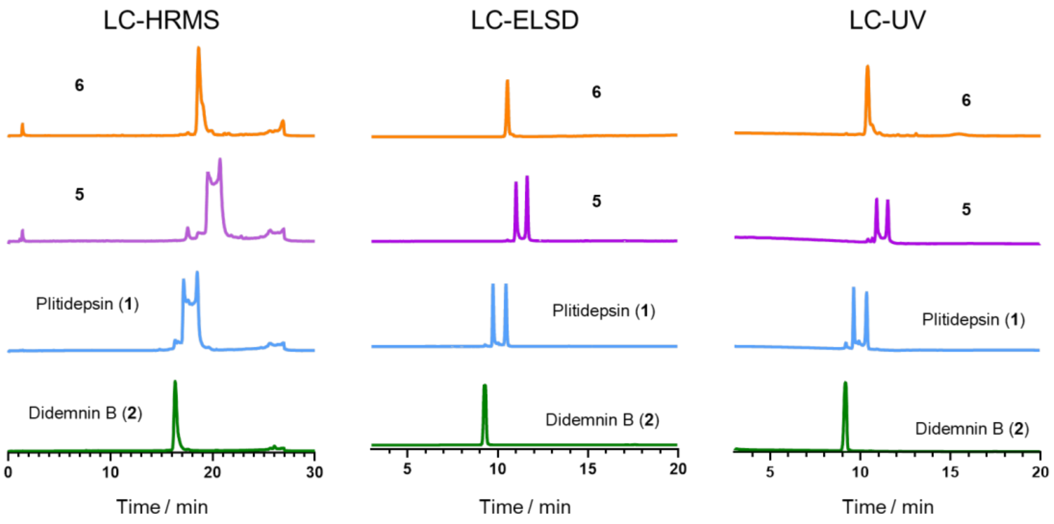


**Figure S43** Didemnin B (**2**), plitidepsin (**1**), compound **5** and **6** inhibit HSV-1-GFP (0.01 MOI) replication in Vero-E6 cells. Viral replication was monitored by IncuCyte S3/SX1 in the presence of the indicated drugs at a concentration of 100 nM. Representative pictures with green fluorescence plus cells or green fluorescence only are shown.

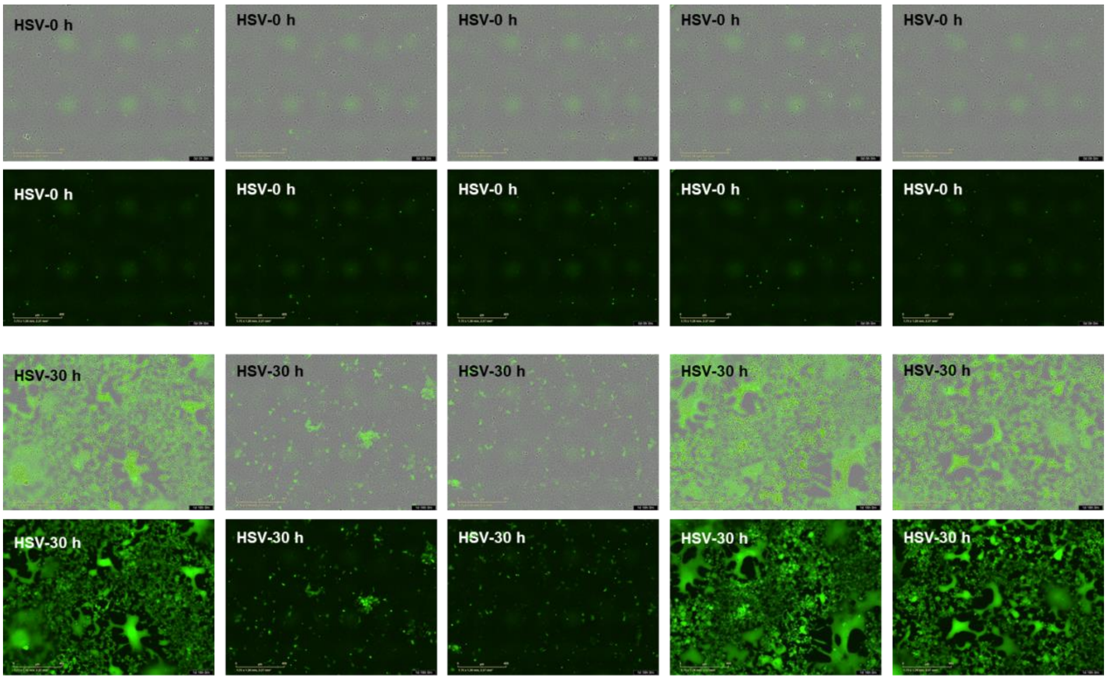


Virus only | Didemnin B (**2**) | Plitidepsin (**1**) | **5** | **6**

**Figure S44** Didemnin B (**2**), plitidepsin (**1**), compound **5** and **6** inhibit VSV-GFP (0.01 MOI) replication in Vero-E6 cells. Viral replication was monitored by IncuCyte S3/SX1 in the presence of the indicated drugs at a concentration of 100 nM. Representative pictures with green fluorescence plus cells or green fluorescence only are shown.

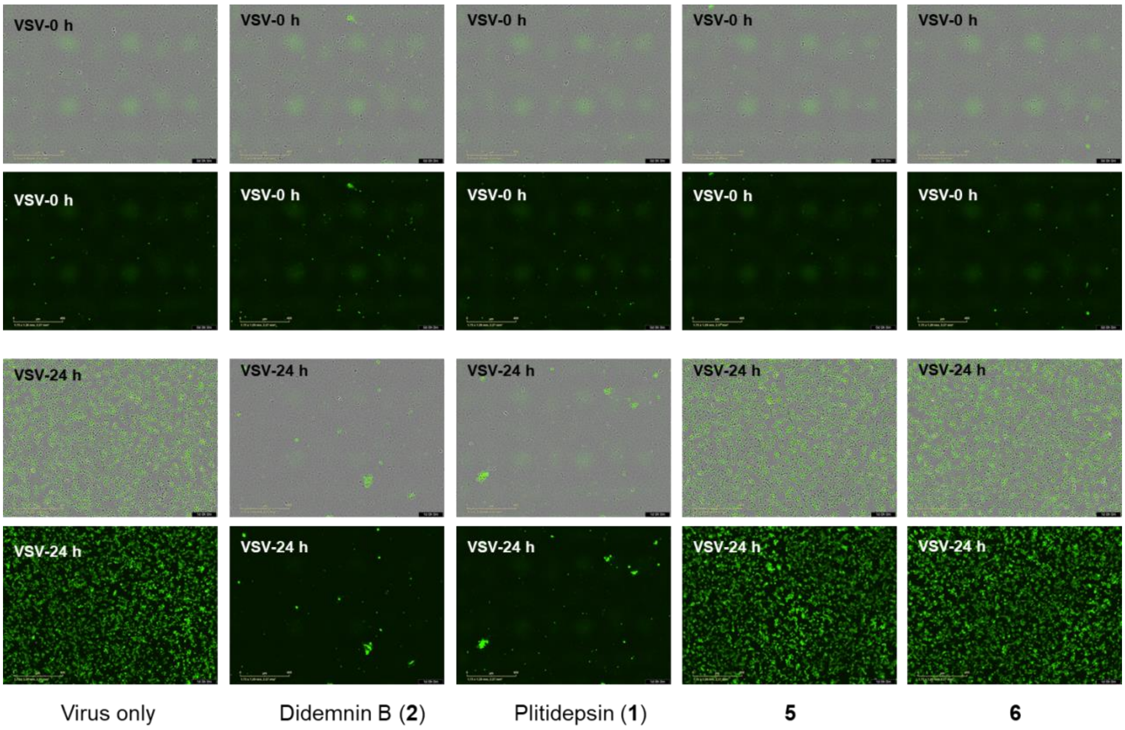
